# Non-invasive Analysis of Peptidoglycan from Living Animals

**DOI:** 10.1101/2023.07.21.549941

**Authors:** Karl L. Ocius, Sree H. Kolli, Saadman S. Ahmad, Jules M. Dressler, Mahendra D. Chordia, Brandon L. Jutras, Melanie R. Rutkowski, Marcos M. Pires

## Abstract

The role of the intestinal microbiota in host health is increasingly revealed in its contributions to disease states. The host-microbiome interaction is multifactorial and dynamic. One of the factors that has recently been strongly associated with host physiological responses is peptidoglycan from bacterial cell walls. Peptidoglycan from gut commensal bacteria activate peptidoglycan sensors in human cells, including the Nucleotide-binding oligomerization domain containing protein 2 (NOD2). When present in the gastrointestinal tract, both the polymeric form (sacculi) and de-polymerized fragments can modulate host physiology, including checkpoint anticancer therapy efficacy, body temperature and appetite, and postnatal growth. To leverage this growing area of biology towards therapeutic prescriptions, it will be critical to directly analyze a key feature of the host-microbiome interaction from living hosts in a reproducible and non-invasive way. Here we show that metabolically labeled peptidoglycan/sacculi can be readily isolated from fecal samples collected from both mice and humans. Analysis of fecal samples provided a non-invasive route to probe the gut commensal community including the metabolic synchronicity with the host circadian clock. Together, these results pave the way for non-invasive diagnostic tools to interrogate the causal nature of peptidoglycan in host health and disease.

## Introduction

The human gastrointestinal (GI) tract is populated by a community of microorganisms that are purported to be involved in a range of biological functions, such as synthesizing vitamins, training the immune system, and protecting the host against pathogens.^1-5^ Disruptions to the human microbiota have been linked to a range of health dysfunctions, including digestive disorders, autoimmune diseases, and obesity. This complex relationship between the microbiota and the host is mediated by multiple factors including the exchange of biologically active molecules. Bacterial products interact with the host immune cells in the gut and influence the development and function of the immune system, which can impact host susceptibility to infections, response to pathogenicity (including cancer), and the development of autoimmune diseases. Recent evidence has revealed that bacterial cell walls are, in fact, biologically active agents to their host organisms (**Figure 1A**).^6-8^

**Figure 1.**
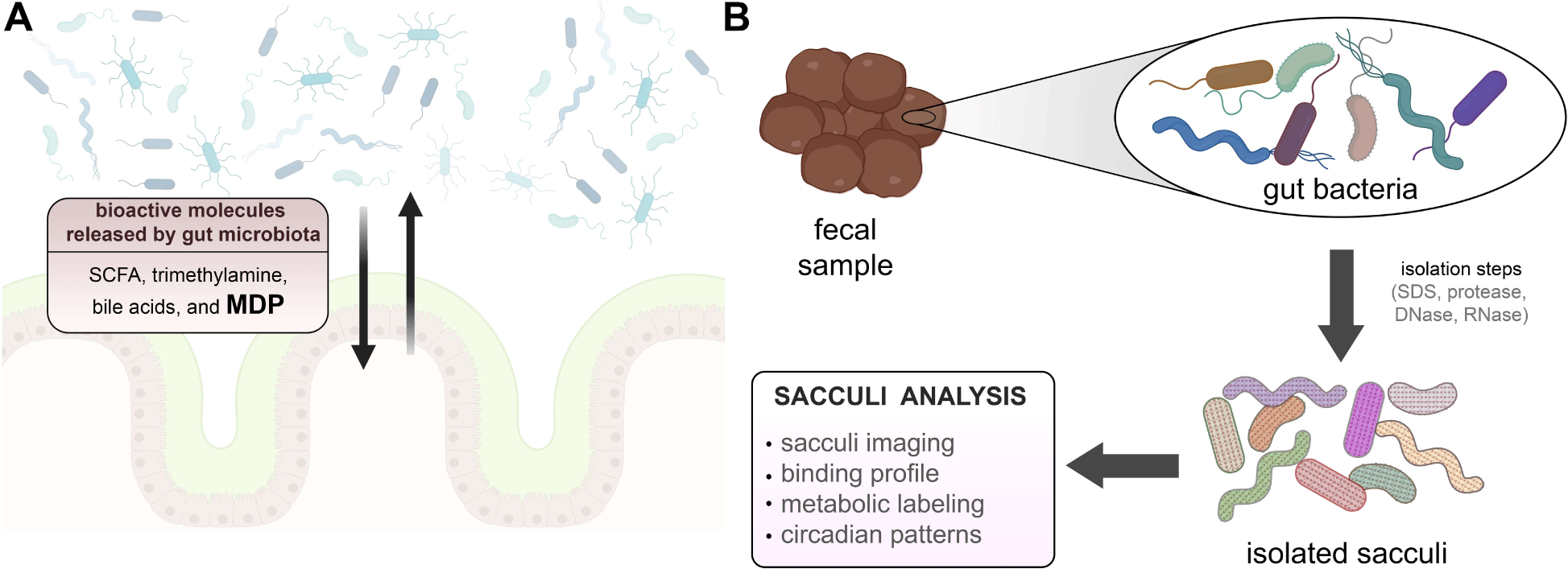
(**A**) Cartoon representation of the interaction between gut microbiota and intestinal lining. This interaction can be driven by the two-way distribution of signaling molecules from and to the host. There are a few molecules that have been identified that are released by gut bacteria, which are known to modulate host physiology. (**B**) Workflow for the analysis of sacculi from fecal samples. Samples are subjected to a series of treatments that ultimately result in the isolation of sacculi that can be analyzed with downstream assays.

Bacterial peptidoglycan is a major component of bacterial cell walls, playing a key role for the defense of bacterial cells.^9,10^ The peptidoglycan scaffold surrounds the entire cell, and this single molecule is known as the bacterial sacculus. Peptidoglycan is composed of unique building blocks that are not present in humans, including a polymeric glycan backbone built from disaccharides of *N*-acetylglucosamine (GlcNAc) and *N*-acetylmuramic acid (MurNAc). A short peptide (the stem peptide) is connected to each MurNAc unit and is crosslinked across neighboring chains forming a mesh-like structure. The stem peptide can vary in composition but is typically 5 amino acids in length, with the sequence L-Ala-*iso*-D-Glu-L-Lys (or *meso*-diaminopimelic acid [*m*-DAP])-D-Ala-D-Ala.^11^ Given the unique chemical composition of peptidoglycan, it represents an ideal biomarker to indicate the presence of bacteria in any system.^10,12^

Organisms have evolved diverse strategies to sense the presence of peptidoglycan as a mode of self-defense and microbiome maintenance.^13,14^ These include peptidoglycan recognition proteins (PGRPs)^15^ and Lysin Motif (LysM) domains that recognize polymeric peptidoglycan,^16,17^ and nucleotide binding and oligomerization domain proteins (NOD1/NOD2) that recognize peptidoglycan fragments.^18-20^ For some receptors, the primary function could be to alert the immune system to the presence of a dangerous bacterial pathogen and trigger an inflammatory host response.^9^ However, for NOD2, recent seminal studies have demonstrated that activation^21^ of NOD2 by agonists, such as peptidoglycan fragments (e.g., muramyl dipeptide, MDP^22-24^), are correlated with positive health outcomes for the host. Peptidoglycan fragments were recently shown to significantly improve responses to checkpoint inhibitors in a cancer model in mice,^25^ modulate body temperature and appetite of mice,^26^ and alleviate Crohn’s disease phenotypes.^8,27^ Critically the health effects were not limited to peptidoglycan fragments; the administration of purified bacterial sacculi from *Lactobacillus plantarum* (*L. plantarum*) led to significant growth improvement in undernourished mice.^6^ Additionally, the impact of sacculi on host biology may not be confined to interactions in the GI tract. Radiolabeled sacculi that were orally administered to mice translocated to the circulatory system and led to subsequent systemic dissemination.^28^

Despite the growing appreciation that gut bacterial sacculi (and its peptidoglycan fragments) are bona fide interspecies signaling molecules,^29^ the direct isolation of sacculi from living organisms has not been properly evaluated. Given the vast diversity of the gut microbiota, including a large fraction that has yet to be cultured *in vitro*, the isolation of sacculi from gut bacterial communities may represent our most reliable access point to polymeric precursors to these peptidoglycan-host signaling molecules. To address the role of sacculi in gut microbiota health, a necessary first step is to directly interrogate sacculi from a living host via a non-invasive and user-friendly approach. Here, we directly isolated sacculi from stool samples to gain insight into the status of the gut microbiome community following downstream analysis including sequencing analysis, multiplexing with binding probes, and metabolic tagging (**Figure 1B**). To date, stool samples have provided the most accessible and least disruptive method to examine the gut microbiome community from a living host. To this end, a large majority of the important studies linking the gut microbiota with various host (human and mice) health impacts have relied on fecal analysis.^30-34^ Our results showed the feasibility of readily isolating sacculi from fecal samples collected from mice and humans; results that create the opportunity to couple sacculi analysis to gut bacterial status. Furthermore, we site selectively tagged peptidoglycan of gut bacteria in live host and recovered them in the fecal samples. We project that the metabolic tags can be leveraged to monitor cell wall biosynthesis/remodeling dynamics in the context of changing external conditions (including light-dark cycles).

## Results

The GI tract has the largest concentration of commensal microorganisms in mammalian hosts such as mice and humans. While analysis by whole genome sequencing from fecal samples has become routine and it affords species level identification of the microbial community, it has been a challenge to leverage this wealth of data towards functional assays. In contrast, to the best of our knowledge, there have not been attempts to isolate and leverage the sacculi of bacteria from stool samples. A unique physicochemical property of sacculi is their resistance to a range of degrative enzymes (proteases, DNAse, RNAses) and detergents; this feature enables the robust and highly reproducible isolation of pure peptidoglycan from cultured cells. We reasoned that this workflow could also be translated to selectively isolate sacculi from the highly complex stool sample matrix. In doing so, we hypothesized that it would be possible to leverage this important biopolymer as a biomarker of the host-microbiota peptidoglycan-signaling axis.

To start, fecal samples from Specific Pathogen Free (SPF) adult mice were collected and subjected to standard sacculi isolation procedures. The sample is initially boiled in Sodium Dodecyl Sulfate (SDS), followed by treatment with trypsin, DNase, and RNAse. We previously showed that sacculi from cells cultured *in vitro* can be readily analyzed by flow cytometry given that their size mirrors that of live bacterial cells.^35^ As expected, the scatter profile of the fecal sacculi sample was similar to that of sacculi from *Lactobacillus casei* (*L. casei*) (**Figure 2A**). *L. casei* is part of the natural flora of humans and can be used as a model organism.^36^ The sacculi sample was then subjected to a number of treatments with agents that have affinity towards the stem peptide or the saccharide backbone on bacterial sacculi to confirm the composition of the isolated biopolymer.

**Figure 2.**
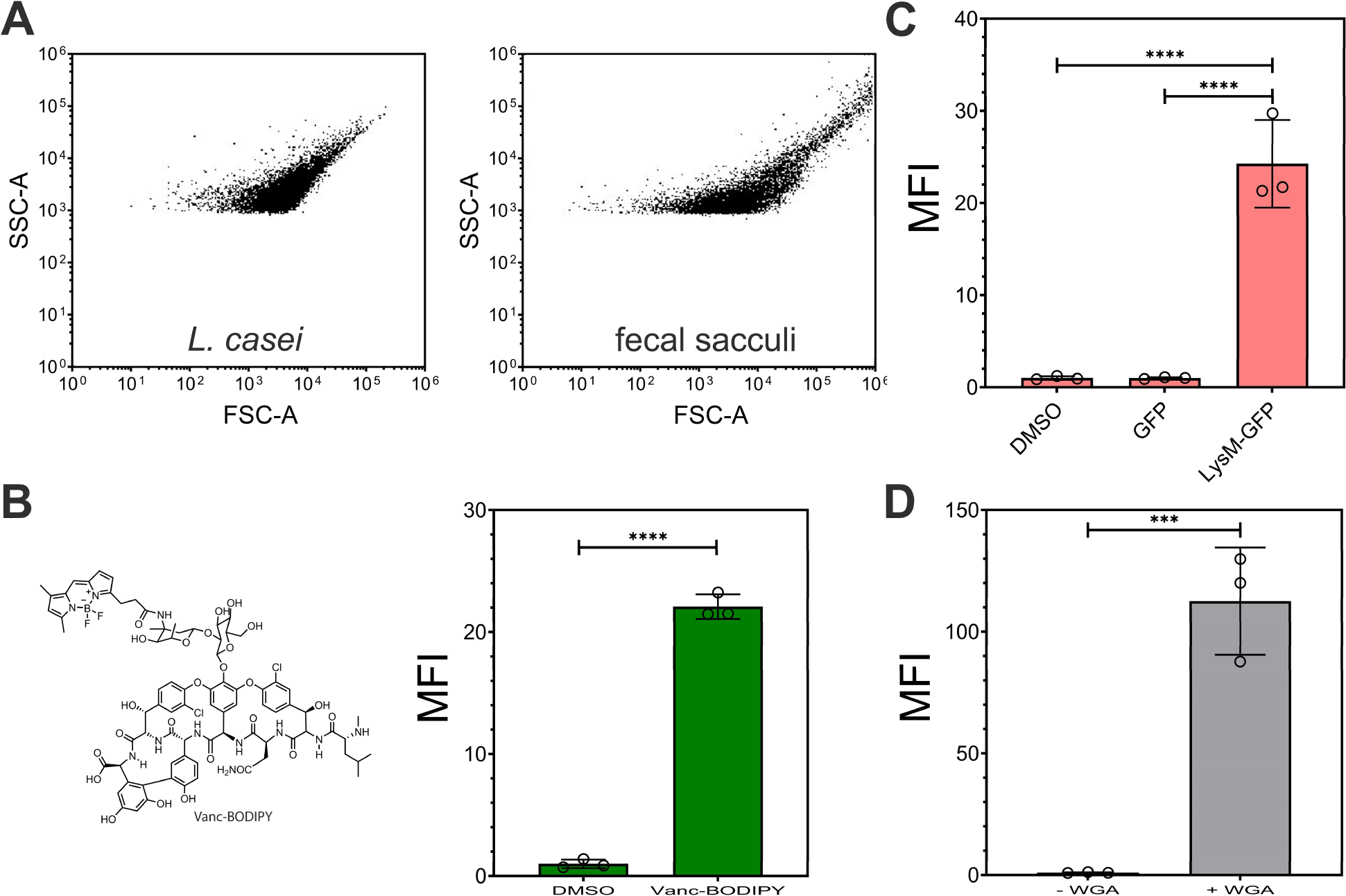
(**A**) Forward and side scatter plots of sacculi isolated from *L. casei* cultured *in vitro* and fecal sacculi from mice. (**B**) Flow cytometry analysis of sacculi isolated from fecal samples of mice in the presence of DMSO or Vanc-BODIPY (2 μg/mL) for 60 mins then washed with PBS. (**C**) Flow cytometry analysis of sacculi isolated from fecal samples of mice in the presence of Lysm-GFP, GFP, or no protein (1 μM) for 60 mins then washed with PBS (**D**) Flow cytometry analysis of sacculi isolated from fecal samples of mice in the presence of DMSO or Fluorescein WGA (1 μM) for 60 mins then washed with PBS. Mean fluorescence intensity (MFI) is the ratio of fluorescence levels above the control (DMSO) treatment from 10000 events. *P*-values were determined by a two-tailed *t*-test (* denotes a *p*-value < 0.05, ** < 0.01, ***<0.001, ns = not significant).

Vancomycin is a glycopeptide that selectively binds to the terminal D-Ala-D-Ala motif on the stem peptide of the peptidoglycan. When linked to the fluorescent dye BODIPY, this conjugate labels whole cells and isolated sacculi.^37^ Incubation of the fecal sacculi sample with Vanc-BODIPY led to a large shift in fluorescence levels, an indication that the material contained the D-Ala-D-Ala motif, which is consistent with isolated sacculi (**Figure 2B**). As expected, a similar profile was also observed using the sacculi from cultured *L. casei* (**Figure S1**). While vancomycin is only biologically active against live Gram-positive bacteria, the sacculi isolation steps will expose D-Ala-D-Ala on sacculi isolated from both Gram-positive and -negative bacteria. To demonstrate that the fluorescent signals were representative of this binding event, sacculi were also treated with a synthetic analog of the stem peptide, L-Lys-D-Ala-D-Ala. A competition experiment was performed by co-treatment of sacculi with Vanc-BODIPY and L-Lys-D-Ala-D-Ala. Accordingly, lower levels of fluorescence were observed upon the addition of L-Lys-D-Ala-D-Ala in a concentration dependent manner (**Figure S2**). These studies provided evidence that the isolated material had binding signatures consistent with the stem peptide of sacculi.

Two additional sacculi binding reagents were tested next to probe the inclusion of the disaccharide backbone within the isolated biopolymers. LysM domains, which are widely found in nature, bind to the GlcNAc saccharide^38^ motif in the backbone of peptidoglycan.^39^ Isolated LysM domains have been shown to bind whole cells and isolated sacculi.^16,40,41^

Treatment of sacculi from fecal samples with LysM fused to GFP led to an increase in fluorescence associated with the sacculi (**Figure 2C**). Treatment with GFP alone and DMSO resulted in near background levels of fluorescence. Similarly, cells were treated with wheat germ agglutinin (WGA) modified with a fluorescein fluorophore. WGA, also known to bind GlcNAc, was recently shown to bind to sacculi of *Borrelia burgdorferi*.^42^ Treatment of the fecal sacculi with fluorescent WGA resulted in a 120-fold increase in fluorescence, which suggests that the material has GlcNAc (**Figure 2D**), similar to that of Lactobacilli cultured *in vitro* (**Figure S3**).

Appreciating that peptidoglycan stem peptides may have glycine residues^43^ and also scars from the trypsin digestion of proteins that are covalently attached to the peptidoglycan (e.g., sortase^44^ anchored or Braun’s protein^45^), we reasoned that we could enzymatically install probes onto the fecal sacculi using Sortase A from *Staphylococcus aureus*. Sortase A (SrtA) is a transpeptidase that covalently anchors endogenous proteins onto the stem peptide of the peptidoglycan scaffold. More specifically, SrtA recognizes the LPXTG (where X is any amino acid) motif to link the third position amino acid on the stem peptide between T and G on the anchored protein. We^35,46,47^ and others^48^ have previously used synthetic analogs of LPXTG conjugated to a fluorophore on the *N*-terminus to metabolically tag isolated bacterial sacculi from pure culture. Here, we incubated fecal sacculi with Fl-LPMTG (fluorescein linked LPMTG, **Figure 3**) in the presence of SrtA and saw a 46-fold increase in fluorescence associated with the sacculi (**Figure 3A**). In the absence of SrtA, basal levels of fluorescence were observed when fecal sacculi was also co-incubated with Fl-LPMTG. Similar fluorescence profiles were also observed with sacculi from *L. casei* (**Figure S4**). The same SrtA fecal sacculi sample was analyzed by confocal microscopy and the sacculi of the fecal sample matched well with prior profile of whole cells extracted from fecal samples of mice (**Figure 3B**).^49^ Together, we showed that we can use bacteria-specific enzymes to covalently tag the sacculi of gut microbiota harvested from fecal sample.

**Figure 3.**
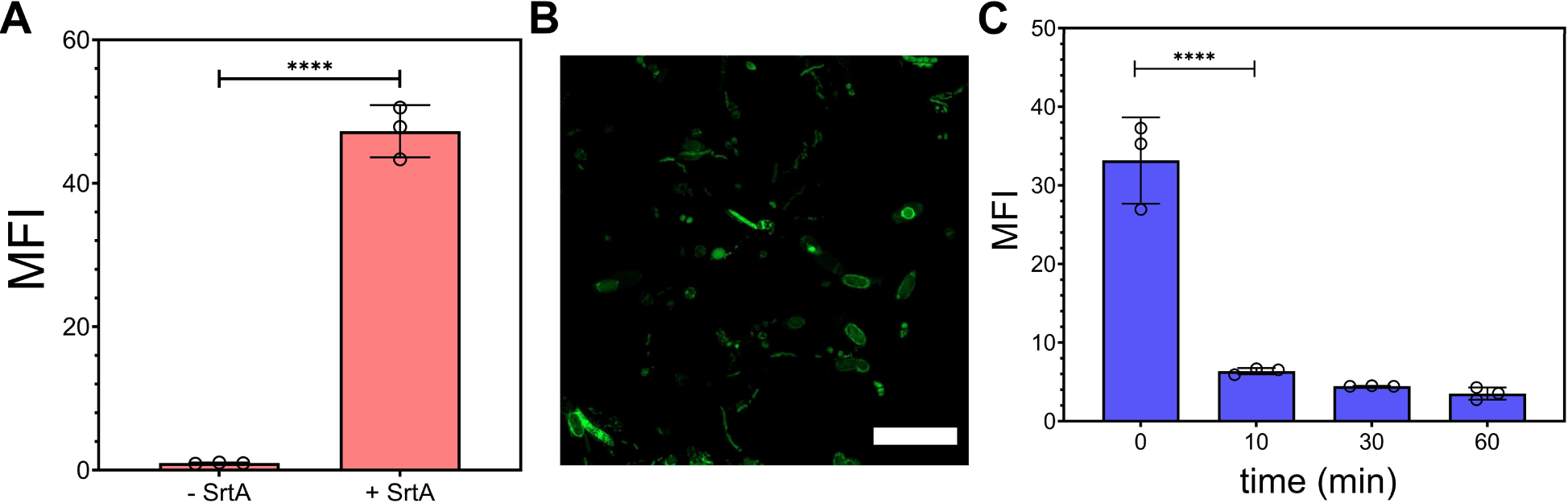
(**A**) Flow cytometry analysis of sacculi isolated from fecal samples of mice in the presence of 100 μM Fl-LPMTG (substrate) and SrtA (20 μM) or without SrtA for 5 h then washed with 100 mM Tris, 5 mM EDTA, pH 7 and freshly added 8 M urea. (**B**) Confocal microscopy of sacculi isolated from fecal samples of mice after tagging with Fl-LPMTG and SrtA; scale bar = 10 μm. (**C**) Flow cytometry analysis of sacculi isolated from fecal samples of mice after tagging with Fl-LPMTG and SrtA in the presence of mutanolysin (25 μg/mL). Samples were monitored across varying time points. Mean fluorescence intensity (MFI) is the ratio of fluorescence levels above the control (-SrtA) treatment from 10000 events. *P*-values were determined by a two-tailed *t*-test (* denotes a *p*-value < 0.05, ** < 0.01, ***<0.001, ****<0.0001, ns = not significant).

Given the polymeric saccharide backbone of peptidoglycan, it may be susceptible to digestion by muramidases such as lysozyme and mutanolysin. *O*-acetylation of muramic acid can result in lysozyme resistance.^50^ Considering the evidence that there is better structural coverage of fecal samples with mutanolysin than lysozyme^51^ in the cell disruption step, and that there is evidence that some gut bacteria include *O*-acetylation^52^, we chose to test muramidase activity with mutanolysin. Sacculi that had been pre-labeled with Fl-LPMTG was incubated with mutanolysin and fluorescence levels were periodically measured. Our data showed that there is a rapid decrease in fluorescence within 10 mins, a clear indication that the isolated material is peptidoglycan in nature (**Figure 3C**). Combined, this broad range of techniques provided foundational evidence that the isolated biopolymer from the fecal samples of mice were primarily sacculi.

Having established sacculi can be readily isolated from the fecal pellets of mice, we sought to demonstrate that a similar workflow of steps should be readily adoptable to human stool samples. In human research, stool samples are conveniently deployed as the primary biospecimen to evaluate the composition and functionality of the human gut microbiota. This is primarily due to the abundance of biomass and the practicality of collection and it has been shown to serve as a valuable proxy for investigating the luminal gut microbiome.^1^ We obtained fecal samples (deidentified) from human donors and matched the steps we had established from mice fecal samples to isolate the bacterial sacculi. Satisfyingly, large increases in fluorescence levels were observed through the treatment of Vanc-BODIPY, WGA, and SrtA (**Figure 4A-C**). These results suggest that the sacculi from fecal samples can be conveniently isolated in the course of approximately 8 h, which provides a potential handle to interrogate the health of a human patient in the event of a pathology that is linked to gut microbiota disturbance. Finally, the isolated sacculi was digested with muramidases and the soluble fragments were analyzed by liquid-chromatography mass spectrometry (**Figure 4D**. Our results showed that a number of the chromatographic peaks had mass (m/z) signatures consistent with muropeptides that included some with a lysine or *m*-DAP in the 3^rd^ position. The remaining peptides could potentially be fragments that were covalently anchored to the sacculi by native transpeptidases. These results furthered confirmed that the isolated insoluble material was enriched in muropeptides.

**Figure 4.**
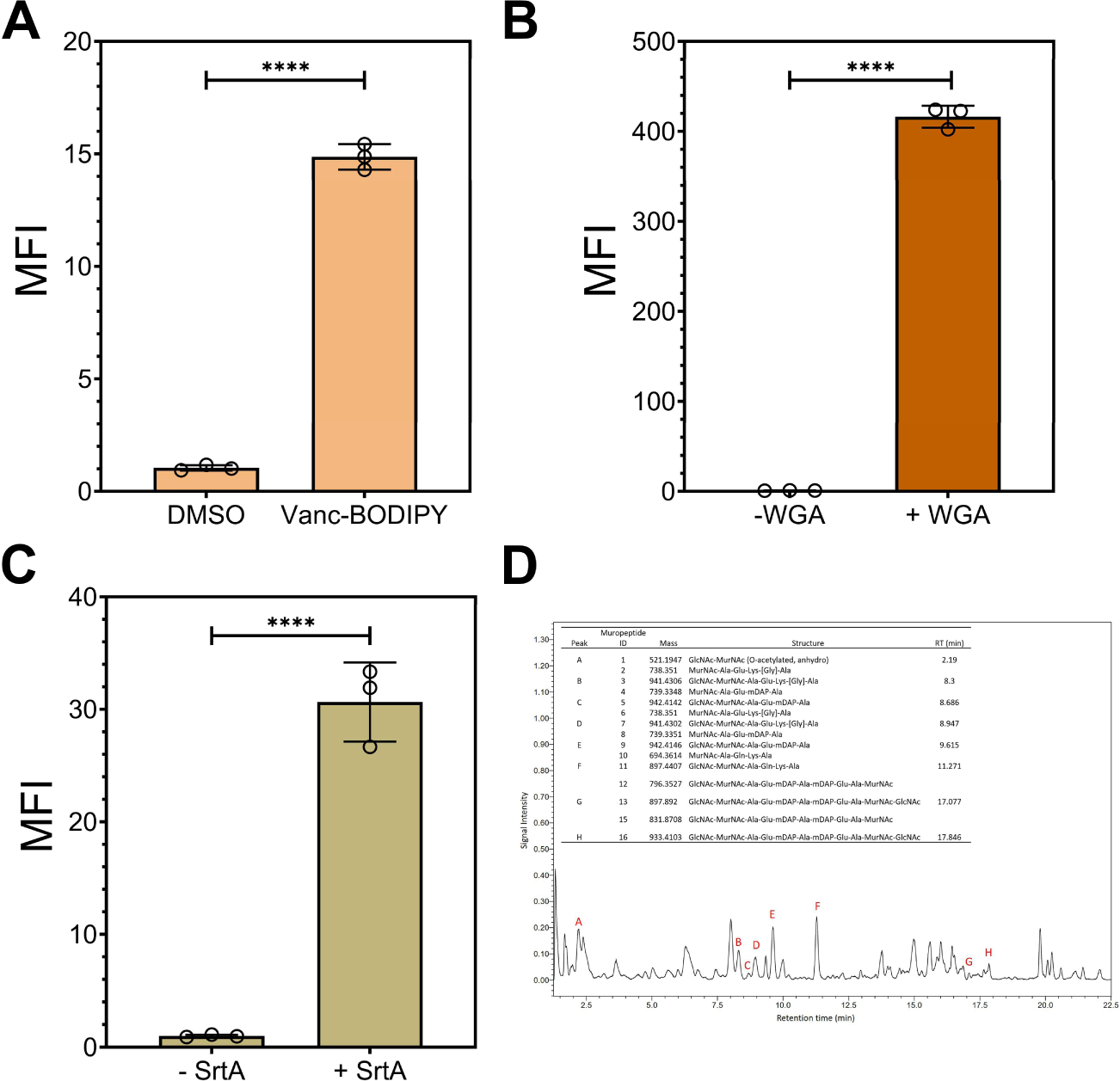
(**A**) Flow cytometry analysis of sacculi isolated from fecal samples of humans in the presence of DMSO or Vanc-BODIPY (2 μg/mL) for 60 mins then washed with PBS. (**B**) Flow cytometry analysis of sacculi isolated from fecal samples of humans in the presence or absence of WGA (1 μM) for 60 mins then washed with PBS. (**C**) Flow cytometry analysis of sacculi isolated from fecal samples of mice in the presence of 100 μM Fl-LPMTG (substrate) and SrtA (20 μM) or without SrtA for 5 h then washed with 100 mM Tris, 5 mM EDTA, pH 7 and freshly added 8 M urea. (**D**) LC-MS chromatogram of processed human fecal samples (black). Muropeptide-containing peaks are labeled (red) and corresponding muropeptide identities (inset table) are shown. MS2 spectra for each peak was scanned for (204.085 *m/z*) and a GlcNAc ring cleavage product (138.053 m/z) and ion masses were compared to known muropeptide masses to assign identity. Data shown represent the results attained from two technical replicates.

We then set out to metabolically label peptidoglycan of gut bacteria in live mice and isolate them directly from fecal samples (**Figure 5**). The goal was to show that the installation of peptidoglycan-specific probes could be detected by a non-invasive method to monitor the metabolism of gut bacteria. Metabolic labeling of the peptidoglycan of cultured bacteria has been heavily explored for studying cell wall dynamics.^53-63^ The simple incubation of synthetic analogs of peptidoglycan with bacterial cells results in their incorporation by promiscuous cell wall enzymes; analogs can be modified with fluorophores or click handles, thus providing routes to illuminating cell wall biosynthesis. In 2017, Hudak et al. (and since then others^64-66^) conjugated a fluorophore to a D-amino acid metabolic tag and found that this probe labeled bacteria in the gut of mice. At the same time, our laboratory showed *in vivo* metabolic labeling of peptidoglycan in *Caenorhabditis elegans*^67^ followed by a recent demonstration in mice.^68^

**Figure 5.**
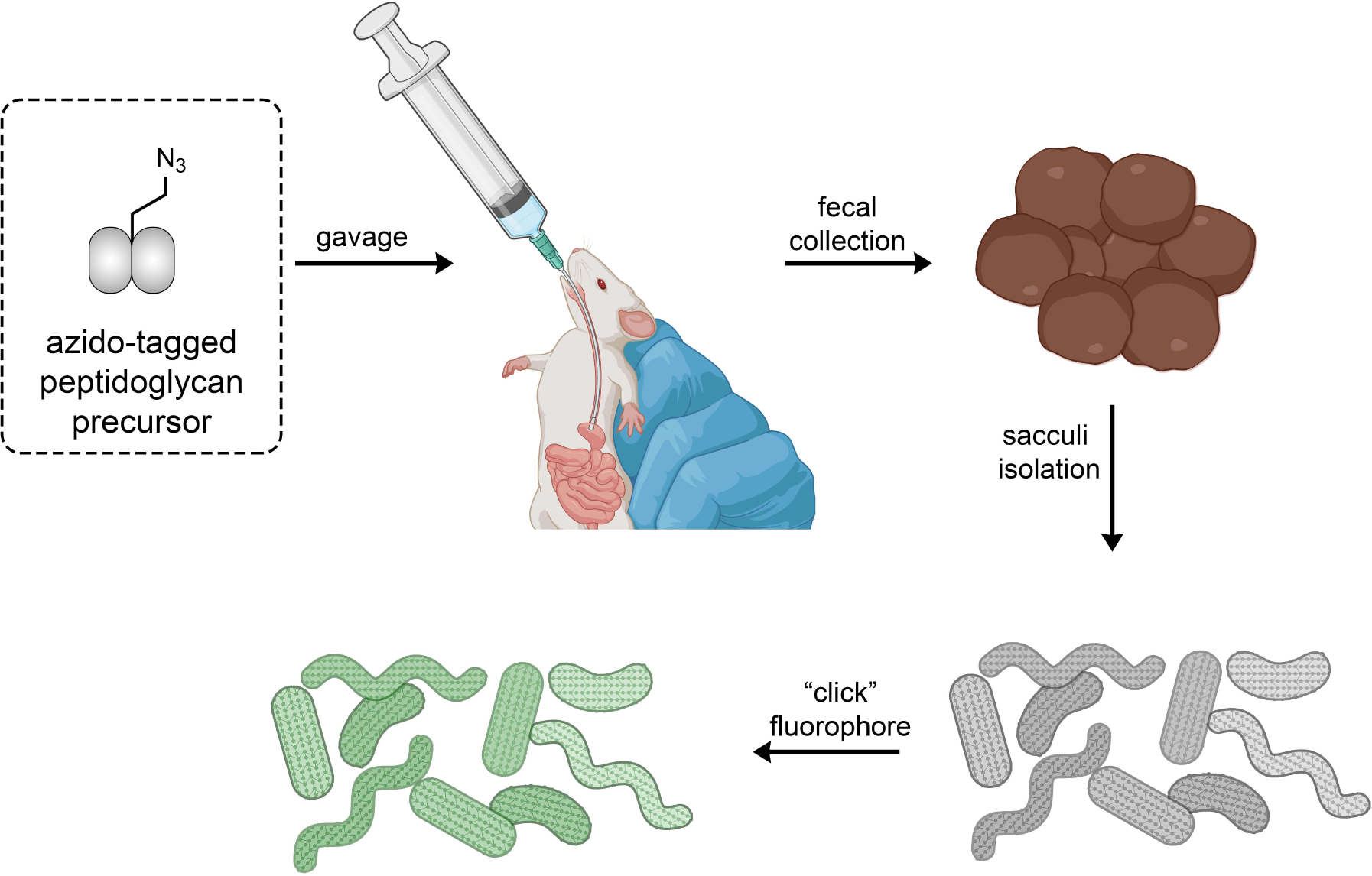
Workflow of the oral administration of a synthetic peptidoglycan analog bearing an azido tag. Following the incorporation of the tag into the cell wall of live bacteria, the fecal samples are collected, and the sacculi are isolated. Finally, a click reaction is performed on the isolated sacculi to install the fluorophore. Sacculi that have been metabolically labeled can subsequently analyzed with various techniques.

To initially benchmark the metabolic labeling conditions, *L. casei* was cultured with DMSO or in the presence of **D-KAz** (**Figure 6A**). **D-KAz** bears an azido tag and it is expected to be metabolically incorporated throughout the peptidoglycan scaffold. Single D-amino acids tags become inserted into the stem peptide of the peptidoglycan by endogenous bacterial transpeptidases, which are responsible for peptidoglycan crosslinking (**Figure 6B**).^69-71^ The installation of the azido group can be revealed with a copper-catalyzed click reaction to an alkyne-modified fluorescein.^72^ *L. casei* cultured in the presence of **D-KAz** led to a 31-fold increase in fluorescence levels relative to untreated cells when cells were analyzed as intact structures (**Figure 6C**). As an alternative method to show the installation of the azido groups on the peptidoglycan, sacculi was first extracted from *L. casei* grown with DMSO or **D-KAz**. The isolated sacculi was then reacted with the fluorophore and a similar level of fluorescence increase was observed. These results indicate that the azido groups found in the whole cells are likely exclusively within the peptidoglycan. More importantly, these findings illustrate the chemical stability of the azido tag to the steps associated with sacculi isolation.

**Figure 6.**
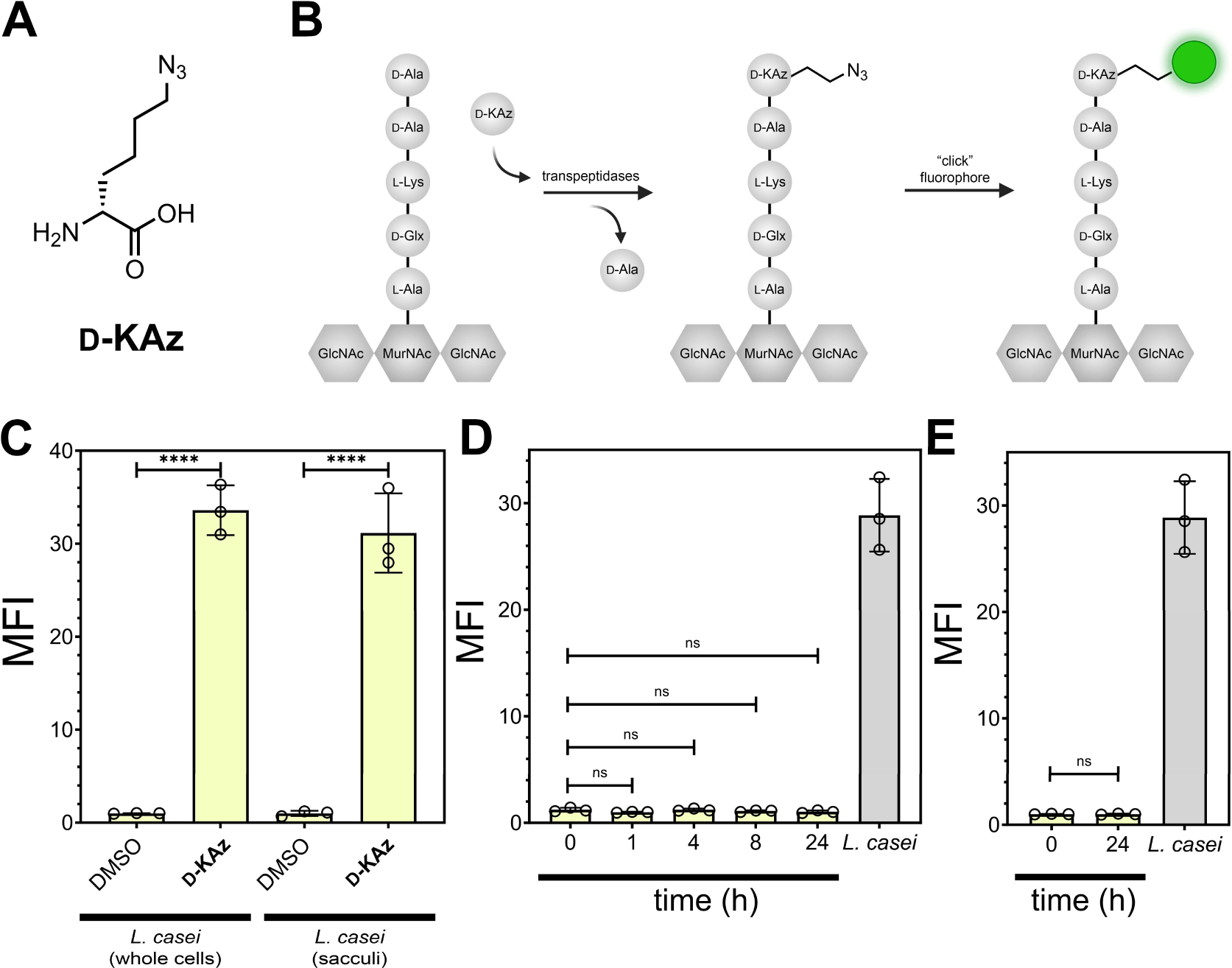
(**A**) Chemical structure of **D-KAz**. (**B**) Schematic cartoon representation of the swapping of the terminal D-alanine for the modified D-amino acid in the media. After installation of the azido tag, a click reaction results in a covalent modification with the fluorophore. (**C**) Flow cytometry analysis of whole cells or sacculi of *L. casei*. *L. casei* cells were treated overnight with 1 mM of **D-KAz** or DMSO then labeled as whole cells by treating with 30 μM of alkyne-fluorescein. Alternatively, cells were subjected to sacculi isolation steps before performing the click reaction. (**D**) Mice were orally dosed with 5 mM of **D-KAz** 2X one hour apart in a daylight cycle. Following the second dosing, the animals were moved to a new cage. Fecal samples were collected at the designated times and subjected to sacculi isolation and a click reaction to install the fluorophore. The levels were compared to sacculi of *L. casei* labeled *in vitro* with **D-KAz**. (**E**) The caecum contents of the mice from (**D**) at time 24 h were harvested, bacterial cells were retrieved, and the sacculi was isolated before performing a click reaction with a fluorophore. The levels were compared to sacculi of *L. casei* labeled *in vitro* with **D-KAz**. Mean fluorescence intensity (MFI) is the ratio of fluorescence levels above the control (DMSO) treatment from 10000 events. *P*-values were determined by a two-tailed *t*-test (* denotes a *p*-value < 0.05, ** < 0.01, ***<0.001, ****<0.0001, ns = not significant).

With the metabolic conditions benchmarked, we orally administered mice with **D-KAz** to analyze labeled sacculi from stool samples. Mice were dosed two times (one hour apart) with **D-KAz** in PBS and subsequently fecal pellets were collected at several time points following the second oral dosing before being subjected to the sacculi isolation steps (**Figure 6D**). No statistically significant differences in sacculi fluorescence levels were found up to 24 hours after dosing of the peptidoglycan tag. Two likely scenarios may have been responsible for these results: the level of metabolic labeling was insufficient under the conditions that were tested and resulted in low levels of azide-tagging, or there had not been enough time for the tagged cells to passage through the gut to the stool sample. To determine if labeled cells had not yet moved to the stool samples, the caecum contents were harvested. Cells found in this region of the GI tract also had sacculi with fluorescence levels near background, suggesting that poor labeling incorporation or retention may have been the reason for lack of fluorescence signals in the sacculi samples (**Figure 6D**).

To increase the overall bacterial cell wall labeling levels, an alternative metabolic tagging strategy was evaluated. Instead of single amino acids, we tested dipeptides that are mimics of the cytosolic peptidoglycan precursor, D-Ala-D-Ala. We^73,74^ and others^58,75^ had previously demonstrated that synthetic D-Ala-D-Ala analogs displaying click chemistry handles (e.g., azide or alkyne) can be installed into peptidoglycan precursors in the intracellular space of bacteria (**Figure 7A**). Most relevant to the proposed change in labeling strategy, we had previously found that labeling levels were generally higher with dipeptide metabolic tags relative to single amino acids.^73,74^ We synthesized **DAzDA** that displays an azido tag on the alanine sidechain of the *N*-terminal amino acid (**Figure 7A**) and tested its incorporation efficiency *in vitro* using *L. casei*. Treatment of cells with **DAzDA** led to a 210-fold increase in fluorescence levels compared to untreated whole cells (**Figure 7B**), which is considerably higher than **D-KAz** labeling levels. As with the single amino acid probe, the azido group was evidently stable through the sacculi isolation steps and the sacculi fluorescence levels from **DAzDA** labeling were similarly higher. Importantly, cellular labeling with **DAzDA** was observed by 30 minutes after incubation, thus establishing that short incubation times can be sufficient for peptidoglycan incorporation (**Figure S5**). Finally, confocal microscopy analysis showed that the labeling pattern observed with **DAzDA** was consistent with the expected sacculi structure (**Figure S6**).

**Figure 7.**
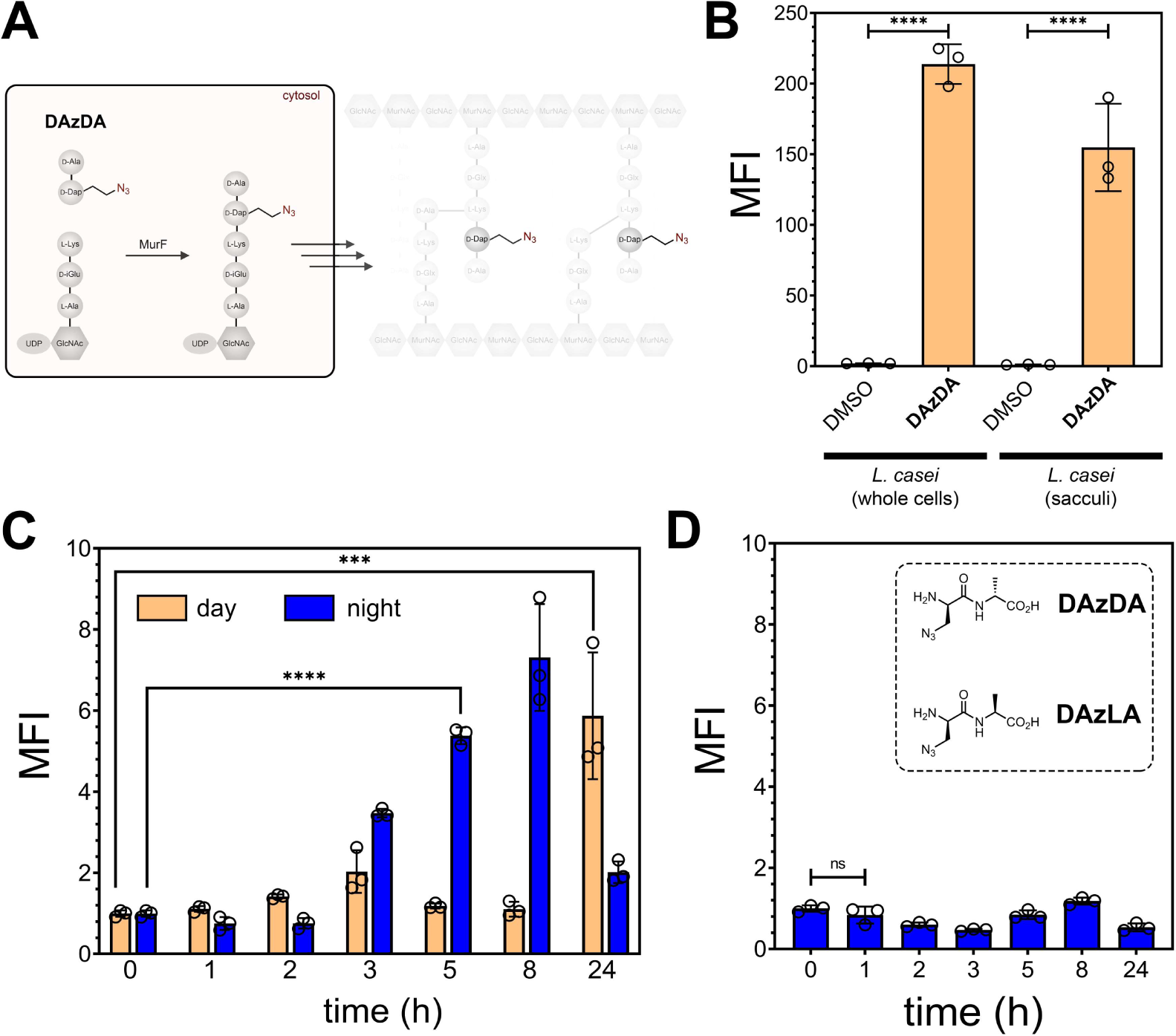
(**A**) Schematic cartoon showing how synthetic analogs of D-Ala-D-Ala enter the biosynthetic pathway at the MurF ligation step. (**B**) Flow cytometry analysis of whole cells or sacculi of *L. casei*. *L. casei* cells were treated overnight with 1 mM of **DAzDA** or DMSO then labeled as whole cells by treating with 30 μM of alkyne-fluorescein. Alternatively, cells were subjected to sacculi isolation steps before performing the click reaction. (**C**) SPF mice were orally dosed with 5 mM of **DAzDA** 2X one hour apart. Following the second dosing, the animals were moved to a new clean cage. Fecal samples were collected at the designated times and subjected to sacculi isolation and a click reaction to install the fluorophore. The process was performed during daylight and during the night cycles. (**D**) SPF mice were orally dosed with 5mM of **DAzLA** 2X one hour apart. Following the second dosing, the animals were moved to a new clean cage. Fecal samples were collected at the designated times and subjected to sacculi isolation and a click reaction to install the fluorophore. The process was performed during the night cycle. Mean fluorescence intensity (MFI) is the ratio of fluorescence levels above the control (DMSO) treatment from 10000 events. *P*-values were determined by a two-tailed *t*-test (* denotes a *p*-value < 0.05, ** < 0.01, ***<0.001, ****<0.0001, ns = not significant).

Mice were next dosed with **DAzDA** to probe for improved metabolic tagging of sacculi in live mice. As before, mice were dosed with the metabolic tag twice (one hour apart) and fecal samples were collected following the second administration. In the first round, mice were dosed during the day light cycle starting early in the morning. Interestingly, the only large difference was between time 0 and time 24 h, without significant change in any of the intervening time points (**Figure 7C**). We wondered whether the circadian rhythm of the host could be impacting the incorporation of the metabolic tag. After all, the incorporation of the metabolic tag is tightly linked with the overall metabolic processing of the bacterial cell wall. Moreover, it had been shown that the gut microbiome exhibits diurnal variations in its composition.^76,77^ Certain microbial species and their metabolic activities display rhythmic patterns that are influenced by the circadian clock of the host.

For example, the abundance and diversity of specific bacteria in the gut can vary throughout the day. To test the possibility that metabolic labeling of the sacculi levels could be impacted by circadian clocks, **DAzDA** was dosed at night and fecal samples were collected following the last administration as the day experiment. A distinctly different pattern of sacculi metabolic labeling emerged (**Figure 7C**). A steady increase in sacculi fluorescence levels was observed throughout the night and 5 h following administration, there was a marked increase in sacculi relative to the initial measurements. By 24 h, the fluorescence levels were approaching that of initial measurements. Confocal analysis of samples isolated from *in situ* labeled bacteria yielded structures consistent with sacculi (**Figure S7**).

To test the specificity of the metabolic labeling, a new dipeptide was synthesized that was a diastereomeric analog that contained an L-amino acid on the *C*-terminus (**Figure 7C**). We^73,74^ previously showed that the stereocenter of D-Ala-D-Ala analogs is crucial for recognition by the biosynthetic machinery. Satisfyingly, mice dosed with **DAzLA** resulted in sacculi with near background fluorescence levels throughout the time sampling time window. These results are consistent with a bacterial-specific incorporation of **DAzLA** in live bacteria cells in the mice. We believe that metabolic labeling of gut microbiota can be a tool to study the metabolic kinetics of gut bacteria in a non-invasive and -disruptive method that can be repeatedly probed over time using stool samples.

## Conclusion

In conclusion, we described the non-invasive analysis and metabolic labeling of bacterial peptidoglycan in the gut microbiota. Fecal samples from mice were subjected to sacculi isolation procedures, and various techniques were employed to examine and label the peptidoglycan. The experiments revealed that the isolated sacculi displayed similar characteristics to those of known bacterial sacculi, confirming their identity. Multiple binding reagents, including vancomycin, LysM domains, and WGA, were used to detect specific components of the peptidoglycan structure and confirm successful sacculi isolation. Additionally, we utilized Sortase A to enzymatically install probes onto the fecal sacculi, and fluorescence assays demonstrated successful incorporation. Metabolic labeling of peptidoglycan in live mice was also demonstrated using synthetic peptidoglycan analogs. The results showed specific tagging of sacculi, indicating successful incorporation of the analogs into the peptidoglycan scaffold. Interestingly, the time of dosing influenced the metabolic labeling levels, suggesting a potential impact of the host’s circadian rhythm on bacterial cell wall biosynthesis. Additionally, we demonstrated the feasibility of non-invasive and repeated probing of gut bacterial metabolism using metabolic labeling techniques, providing a potential tool for studying the dynamics of gut microbiota over time. We propose that directly isolating sacculi from stool samples can offer a novel and biologically relevant assessment route to the interaction between gut microbiota and the host.

## Acknowledgement

This study was supported by the NIH grant R01AI181139 (KLO, MDC, MMP). Partial support for both SSA and JMD was provided by the NIH (R21AI159800, R01AI173256, R01AI178711) and the Global Lyme Alliance, all awarded to BLJ and MMP.

## Supporting Information

Additional figures, tables, and materials/methods are included in the supporting information file.

## References

1 Eckburg, P. B. et al. Diversity of the human intestinal microbial flora. Science 308, 1635–1638 (2005). 10.1126/science.1110591

2 Round, J. L. & Mazmanian, S. K. The gut microbiota shapes intestinal immune responses during health and disease. Nat Rev Immunol 9, 313–323 (2009). 10.1038/nri2515

3 Qin, J. et al. A human gut microbial gene catalogue established by metagenomic sequencing. Nature 464, 59–65 (2010). 10.1038/nature08821

4 Human Microbiome Project, C. Structure, function and diversity of the healthy human microbiome. Nature 486, 207–214 (2012). 10.1038/nature11234

5 Belkaid, Y. & Hand, T. W. Role of the microbiota in immunity and inflammation. Cell 157, 121–141 (2014). 10.1016/j.cell.2014.03.011

6 Schwarzer, M. et al. Microbe-mediated intestinal NOD2 stimulation improves linear growth of undernourished infant mice. Science 379, 826–833 (2023). 10.1126/science.ade9767

7 Royet, J., Gupta, D. & Dziarski, R. Peptidoglycan recognition proteins: modulators of the microbiome and inflammation. Nat Rev Immunol 11, 837–851 (2011). 10.1038/nri3089

8 Gao, J. et al. Gut microbial DL-endopeptidase alleviates Crohn’s disease via the NOD2 pathway. Cell Host Microbe 30, 1435–1449 e1439 (2022). 10.1016/j.chom.2022.08.002

9 Royet, J. & Dziarski, R. Peptidoglycan recognition proteins: pleiotropic sensors and effectors of antimicrobial defences. Nat Rev Microbiol 5, 264–277 (2007). 10.1038/nrmicro1620

10 Wolf, A. J. & Underhill, D. M. Peptidoglycan recognition by the innate immune system. Nat Rev Immunol 18, 243–254 (2018). 10.1038/nri.2017.136

11 Egan, A. J. F., Errington, J. & Vollmer, W. Regulation of peptidoglycan synthesis and remodelling. Nat Rev Microbiol 18, 446–460 (2020). 10.1038/s41579-020-0366-3

12 Guan, R. & Mariuzza, R. A. Peptidoglycan recognition proteins of the innate immune system. Trends Microbiol 15, 127–134 (2007). 10.1016/j.tim.2007.01.006

13 Dworkin, J. The medium is the message: interspecies and interkingdom signaling by peptidoglycan and related bacterial glycans. Annu Rev Microbiol 68, 137–154 (2014). 10.1146/annurev-micro-091213-112844

14 Hayes, C. L. et al. Commensal microbiota induces colonic barrier structure and functions that contribute to homeostasis. Sci Rep 8, 14184 (2018). 10.1038/s41598-018-32366-6

15 Dziarski, R. Peptidoglycan recognition proteins (PGRPs). Mol Immunol 40, 877–886 (2004). 10.1016/j.molimm.2003.10.011

16 Yokoyama, C. C. et al. LysMD3 is a type II membrane protein without an in vivo role in the response to a range of pathogens. J Biol Chem 293, 6022–6038 (2018). 10.1074/jbc.RA117.001246

17 He, X. et al. LYSMD3: A mammalian pattern recognition receptor for chitin. Cell Rep 36, 109392 (2021). 10.1016/j.celrep.2021.109392

18 Inohara, Chamaillard, McDonald, C. & Nunez, G. NOD-LRR proteins: role in host-microbial interactions and inflammatory disease. Annu Rev Biochem 74, 355–383 (2005). 10.1146/annurev.biochem.74.082803.133347

19 Fritz, J. H., Ferrero, R. L., Philpott, D. J. & Girardin, S. E. Nod-like proteins in immunity, inflammation and disease. Nat Immunol 7, 1250–1257 (2006). 10.1038/ni1412

20 Travassos, L. H. et al. Nod1 and Nod2 direct autophagy by recruiting ATG16L1 to the plasma membrane at the site of bacterial entry. Nat Immunol 11, 55–62 (2010). 10.1038/ni.1823

21 Lauro, M. L., D’Ambrosio, E. A., Bahnson, B. J. & Grimes, C. L. Molecular Recognition of Muramyl Dipeptide Occurs in the Leucine-rich Repeat Domain of Nod2. ACS Infect Dis 3, 264–270 (2017). 10.1021/acsinfecdis.6b00154

22 Girardin, S. E. et al. Nod2 is a general sensor of peptidoglycan through muramyl dipeptide (MDP) detection. J Biol Chem 278, 8869–8872 (2003). 10.1074/jbc.C200651200

23 Girardin, S. E. et al. Peptidoglycan molecular requirements allowing detection by Nod1 and Nod2. J Biol Chem 278, 41702–41708 (2003). 10.1074/jbc.M307198200

24 Grimes, C. L., Ariyananda Lde, Z., Melnyk, J. E. & O’Shea, E. K. The innate immune protein Nod2 binds directly to MDP, a bacterial cell wall fragment. J Am Chem Soc 134, 13535–13537 (2012). 10.1021/ja303883c

25 Griffin, M. E. et al. Enterococcus peptidoglycan remodeling promotes checkpoint inhibitor cancer immunotherapy. Science 373, 1040–1046 (2021). 10.1126/science.abc9113

26 Gabanyi, I. et al. Bacterial sensing via neuronal Nod2 regulates appetite and body temperature. Science 376, eabj3986 (2022). 10.1126/science.abj3986

27 Lauro, M. L., Burch, J. M. & Grimes, C. L. The effect of NOD2 on the microbiota in Crohn’s disease. Curr Opin Biotechnol 40, 97–102 (2016). 10.1016/j.copbio.2016.02.028

28 Wheeler, R. et al. Microbiota-induced active translocation of peptidoglycan across the intestinal barrier dictates its within-host dissemination. Proc Natl Acad Sci U S A 120, e2209936120 (2023). 10.1073/pnas.2209936120

29 Irazoki, O., Hernandez, S. B. & Cava, F. Peptidoglycan Muropeptides: Release, Perception, and Functions as Signaling Molecules. Front Microbiol 10, 500 (2019). 10.3389/fmicb.2019.00500

30 Gill, S. R. et al. Metagenomic analysis of the human distal gut microbiome. Science 312, 1355–1359 (2006). 10.1126/science.1124234

31 Wu, G. D. et al. Linking long-term dietary patterns with gut microbial enterotypes. Science 334, 105–108 (2011). 10.1126/science.1208344

32 Yatsunenko, T. et al. Human gut microbiome viewed across age and geography. Nature 486, 222–227 (2012). 10.1038/nature11053

33 Goodrich, J. K. et al. Human genetics shape the gut microbiome. Cell 159, 789–799 (2014). 10.1016/j.cell.2014.09.053

34 Turnbaugh, P. J. et al. The effect of diet on the human gut microbiome: a metagenomic analysis in humanized gnotobiotic mice. Sci Transl Med 1, 6ra14 (2009). 10.1126/scitranslmed.3000322

35 Apostolos, A. J., Ferraro, N. J., Dalesandro, B. E. & Pires, M. M. SaccuFlow: A High-Throughput Analysis Platform to Investigate Bacterial Cell Wall Interactions. ACS Infect Dis 7, 2483–2491 (2021). 10.1021/acsinfecdis.1c00255

36 Hill, D. et al. The Lactobacillus casei Group: History and Health Related Applications. Front Microbiol 9, 2107 (2018). 10.3389/fmicb.2018.02107

37 DeDent, A. C., McAdow, M. & Schneewind, O. Distribution of protein A on the surface of Staphylococcus aureus. J Bacteriol 189, 4473–4484 (2007). 10.1128/JB.00227-07

38 Mesnage, S. et al. Molecular basis for bacterial peptidoglycan recognition by LysM domains. Nat Commun 5, 4269 (2014). 10.1038/ncomms5269

39 Rabinovich, G. A. & Toscano, M. A. Turning ‘sweet’ on immunity: galectin-glycan interactions in immune tolerance and inflammation. Nat Rev Immunol 9, 338–352 (2009). 10.1038/nri2536

40 Frankel, M. B. & Schneewind, O. Determinants of murein hydrolase targeting to cross-wall of Staphylococcus aureus peptidoglycan. J Biol Chem 287, 10460–10471 (2012). 10.1074/jbc.M111.336404

41 Snee, M. et al. Peptidoglycan recognition in Drosophila is mediated by LysMD3/4. J Biol Chem, 104758 (2023). 10.1016/j.jbc.2023.104758

42 Davis, M. M. et al. The peptidoglycan-associated protein NapA plays an important role in the envelope integrity and in the pathogenesis of the lyme disease spirochete. PLoS Pathog 17, e1009546 (2021). 10.1371/journal.ppat.1009546

43. https://www.biorxiv.org/content/10.1101/2023.04.17.537164v1.full.pdf.

44 Hendrickx, A. P., Budzik, J. M., Oh, S. Y. & Schneewind, O. Architects at the bacterial surface - sortases and the assembly of pili with isopeptide bonds. Nat Rev Microbiol 9, 166–176 (2011). 10.1038/nrmicro2520

45 Dramsi, S., Magnet, S., Davison, S. & Arthur, M. Covalent attachment of proteins to peptidoglycan. FEMS Microbiol Rev 32, 307–320 (2008). 10.1111/j.1574-6976.2008.00102.x

46 Sabulski, M. J., Pidgeon, S. E. & Pires, M. M. Immuno-targeting of Staphylococcus aureus via surface remodeling complexes. Chem Sci 8, 6804–6809 (2017). 10.1039/c7sc02721d

47 Apostolos, A. J., Kelly, J. J., Ongwae, G. M. & Pires, M. M. Structure Activity Relationship of the Stem Peptide in Sortase A Mediated Ligation from Staphylococcus aureus. Chembiochem 23, e202200412 (2022). 10.1002/cbic.202200412

48 Nelson, J. W. et al. A biosynthetic strategy for re-engineering the Staphylococcus aureus cell wall with non-native small molecules. ACS Chem Biol 5, 1147–1155 (2010). 10.1021/cb100195d

49 Hudak, J. E., Alvarez, D., Skelly, A., von Andrian, U. H. & Kasper, D. L. Illuminating vital surface molecules of symbionts in health and disease. Nat Microbiol 2, 17099 (2017). 10.1038/nmicrobiol.2017.99

50 Davis, K. M. & Weiser, J. N. Modifications to the peptidoglycan backbone help bacteria to establish infection. Infect Immun 79, 562–570 (2011). 10.1128/IAI.00651-10

51 Yuan, S., Cohen, D. B., Ravel, J., Abdo, Z. & Forney, L. J. Evaluation of methods for the extraction and purification of DNA from the human microbiome. PLoS One 7, e33865 (2012). 10.1371/journal.pone.0033865

52 Y., H., et al. High-throughput Automated Muropeptide Analysis (HAMA) Reveals Peptidoglycan Composition of Gut Microbial Cell Walls. Elife eLife.88491.1 (2023).

53 Kuru, E. et al. In Situ probing of newly synthesized peptidoglycan in live bacteria with fluorescent D-amino acids. Angew Chem Int Ed Engl 51, 12519–12523 (2012). 10.1002/anie.201206749

54 Siegrist, M. S. et al. (D)-Amino acid chemical reporters reveal peptidoglycan dynamics of an intracellular pathogen. ACS Chem Biol 8, 500–505 (2013). 10.1021/cb3004995

55 Bisson-Filho, A. W. et al. Treadmilling by FtsZ filaments drives peptidoglycan synthesis and bacterial cell division. Science 355, 739–743 (2017). 10.1126/science.aak9973 355/6326/739 [pii]

56 Pilhofer, M. et al. Discovery of chlamydial peptidoglycan reveals bacteria with murein sacculi but without FtsZ. Nat Commun 4, 2856 (2013). 10.1038/ncomms3856 ncomms3856 [pii]

57 Fleurie, A. et al. MapZ marks the division sites and positions FtsZ rings in Streptococcus pneumoniae. Nature 516, 259–262 (2014). 10.1038/nature13966 nature13966 [pii]

58 Liechti, G. W. et al. A new metabolic cell-wall labelling method reveals peptidoglycan in Chlamydia trachomatis. Nature 506, 507–510 (2014). 10.1038/nature12892

59 Faure, L. M. et al. The mechanism of force transmission at bacterial focal adhesion complexes. Nature 539, 530–535 (2016). 10.1038/nature20121 nature20121 [pii]

60 Qiao, Y. et al. Detection of lipid-linked peptidoglycan precursors by exploiting an unexpected transpeptidase reaction. J Am Chem Soc 136, 14678–14681 (2014). 10.1021/ja508147s

61 Lebar, M. D. et al. Reconstitution of peptidoglycan cross-linking leads to improved fluorescent probes of cell wall synthesis. J Am Chem Soc 136, 10874–10877 (2014). 10.1021/ja505668f

62 Shieh, P., Siegrist, M. S., Cullen, A. J. & Bertozzi, C. R. Imaging bacterial peptidoglycan with near-infrared fluorogenic azide probes. Proc Natl Acad Sci U S A 111, 5456–5461 (2014). 10.1073/pnas.1322727111 1322727111 [pii]

63 Ngo, J. T. et al. Click-EM for imaging metabolically tagged nonprotein biomolecules. Nat Chem Biol 12, 459–465 (2016). 10.1038/nchembio.2076 nchembio.2076 [pii]

64 Wang, W. et al. Assessing the viability of transplanted gut microbiota by sequential tagging with D-amino acid-based metabolic probes. Nat Commun 10, 1317 (2019). 10.1038/s41467-019-09267-x

65 Lin, L. et al. Quantification of Bacterial Metabolic Activities in the Gut by d-Amino Acid-Based In Vivo Labeling. Angew Chem Int Ed Engl 59, 11923–11926 (2020). 10.1002/anie.202004703

66 Wang, W. et al. Metabolic Labeling of Peptidoglycan with NIR-II Dye Enables In Vivo Imaging of Gut Microbiota. Angew Chem Int Ed Engl 59, 2628–2633 (2020). 10.1002/anie.201910555

67 Pidgeon, S. E. & Pires, M. M. Cell Wall Remodeling of Staphylococcus aureus in Live Caenorhabditis elegans. Bioconjug Chem 28, 2310–2315 (2017). 10.1021/acs.bioconjchem.7b00363

68 Apostolos, A. J. et al. Real-time non-invasive fluorescence imaging of gut commensal bacteria to detect dynamic changes in the microbiome of live mice. Cell Chem Biol (2022). 10.1016/j.chembiol.2022.11.010

69 Holtje, J. V. Growth of the stress-bearing and shape-maintaining murein sacculus of Escherichia coli. Microbiol Mol Biol Rev 62, 181–203 (1998). 10.1128/MMBR.62.1.181-203.1998

70 Vollmer, W., Blanot, D. & de Pedro, M. A. Peptidoglycan structure and architecture. FEMS Microbiol Rev 32, 149–167 (2008). 10.1111/j.1574-6976.2007.00094.x FMR094 [pii]

71 Lovering, A. L., Safadi, S. S. & Strynadka, N. C. Structural perspective of peptidoglycan biosynthesis and assembly. Annu Rev Biochem 81, 451–478 (2012). 10.1146/annurev-biochem-061809-112742

72 Kolb, H. C. & Sharpless, K. B. The growing impact of click chemistry on drug discovery. Drug Discov Today 8, 1128–1137 (2003). 10.1016/s1359-6446(03)02933-7

73 Fura, J. M., Pidgeon, S. E., Birabaharan, M. & Pires, M. M. Dipeptide-Based Metabolic Labeling of Bacterial Cells for Endogenous Antibody Recruitment. ACS Infect Dis 2, 302–309 (2016). 10.1021/acsinfecdis.6b00007

74 Sarkar, S., Libby, E. A., Pidgeon, S. E., Dworkin, J. & Pires, M. M. In Vivo Probe of Lipid II-Interacting Proteins. Angew Chem Int Ed Engl 55, 8401–8404 (2016). 10.1002/anie.201603441

75 Garcia-Heredia, A. et al. Peptidoglycan precursor synthesis along the sidewall of pole-growing mycobacteria. Elife 7 (2018). 10.7554/eLife.37243

76 Liang, X., Bushman, F. D. & FitzGerald, G. A. Rhythmicity of the intestinal microbiota is regulated by gender and the host circadian clock. Proc Natl Acad Sci U S A 112, 10479–10484 (2015). 10.1073/pnas.1501305112

77 Thaiss, C. A. et al. Transkingdom control of microbiota diurnal oscillations promotes metabolic homeostasis. Cell 159, 514–529 (2014). 10.1016/j.cell.2014.09.048

