## Supporting Information for "Non-invasive Analysis of Peptidoglycan from Living Animals"

### Table of Contents


| **Supporting Figures** | **S4-S11** |
|---|---|
| **Figure S1:** Labeling of *L. casei* sacculi with VBD | S4 |
| **Figure S2:** Competition of VBD labeled fecal sacculi with L-Lys-D-Ala-D-Ala | S5 |
| **Figure S3:** Labeling of *L. casei* and *L. plantarum* with WGA-Fluorescein | S6 |
| **Figure S4:** Sortase A catalyzed labeling of *L. casei* sacculi | S7 |
| **Figure S5:** Time dependent labeling of *L. casei* sacculi with **DAzDA** | S8 |
| **Figure S6:** Confocal microscopy of **DAzDA** labelled *L. casei* sacculi | S9 |
| **Figure S7:** Click chemistry of gut bacterial sacculi labeled with DAzDA in the absence of the copper catalyst | S10 |
| **Figure S8:** Confocal microscopy of gut bacterial sacculi labelled with **DAzDA** | S11 |
| **Methods and Materials** | **S12-S14** |
| Materials | S12 |
| Mice | S12 |
| Isolation of *L.casei* and *L.plantarum* peptidoglycan/ sacculi | S12 |
| Isolation of gut commensal peptidoglycan/ sacculi from mice and human fecal samples | S12 |
| Vancomycin-BODIPY (VBD) binding and competition Assays | S13 |
| LysM-GFP expression and purification | S13 |
| LysM-GFP and WGA-Fl sacculi binding assay | S13 |
| Sortase A Expression and Purification | S14 |
| Sortase A Enzymatic Assays | S14 |
| Enzymatic degradation of fecal sacculi | S14 |
| Peptidoglycan processing | S14 |
| LC-MS analysis of muramidase digested peptidoglycan | S15 |
| Metabolic labeling of *L. casei* and sacculi isolation | S15 |

| | |
|---|---|
| Time dependent Labeling of *L. casei* sacculi with **DAzDA** | S15 |
| *In-vivo* gut labeling with **DKAz**, **DAzDA,** or **DAzLA** | S15 |
| CuAAC/ click reaction on bacterial peptidoglycan/ sacculi | S16 |
| Confocal imaging of labelled sacculi | S16 |
| **Synthesis and Characterization** | **S17-S23** |
| Scheme S1: **DAzDA and DAzLA** | S17 |
| Scheme S2: **FI-LPMTG** | S18 |
| **DAzDA** and **DAzLA** HRMS and NMR | S19-S21 |
| **FI-LPMTG** analytical HPLC and HRMS | S22-S23 |
| **References** | S24 |

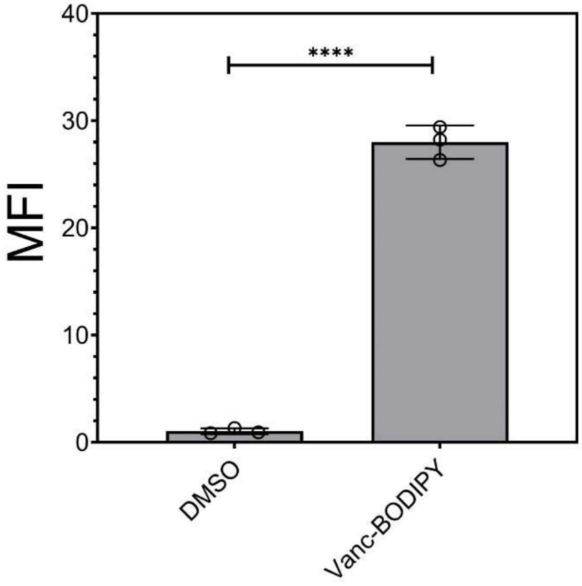


**Figure S1.** Flow cytometry analysis of sacculi isolated from *L. casei* in the presence of Vanc-BODIPY or DMSO (control) for 60 min at 37 °C then washed (3X) with PBS. Mean fluorescence intensity (MFI) is the ratio of fluorescence levels above the control (DMSO) treatment from 10000 events. P-values were determined by a two-tailed t-test (* denotes a p-value < 0.05, ** < 0.01, ***<0.001, ns = not significant).

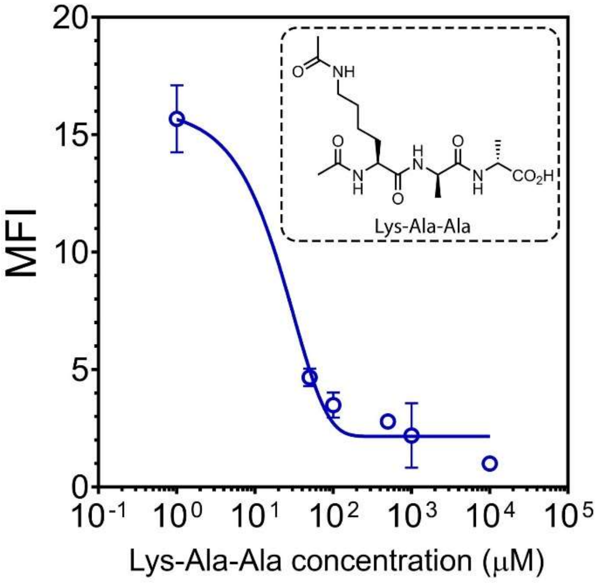


**Figure S2.** Competition of Vanc-BODIPY (VBD) with L-Lys-D-Ala-D-Ala. Sacculi isolated from mouse fecal sample were resuspended in 1X PBS solution containing 2ug/mL VBD or 2 μg/mL VBD in conjunction with either 50, 100, 500, 1000, or 10000 μg/mL of the L-Lys-d-Ala-d-Ala peptide and incubated for 1 h at 37°C, then washed 3 times with 1X PBS, and analyzed by flow cytometry. A concentration dependent decrease in VBD signal was seen with increased concentration of L-Lys-D-Ala-D-Ala peptide. Mean fluorescence intensity (MFI) is the ratio of fluorescence levels above the control (DMSO) treatment from 10000 events. P-values were determined by a two-tailed t-test (* denotes a p-value < 0.05, ** < 0.01, ***<0.001, ns = not significant).

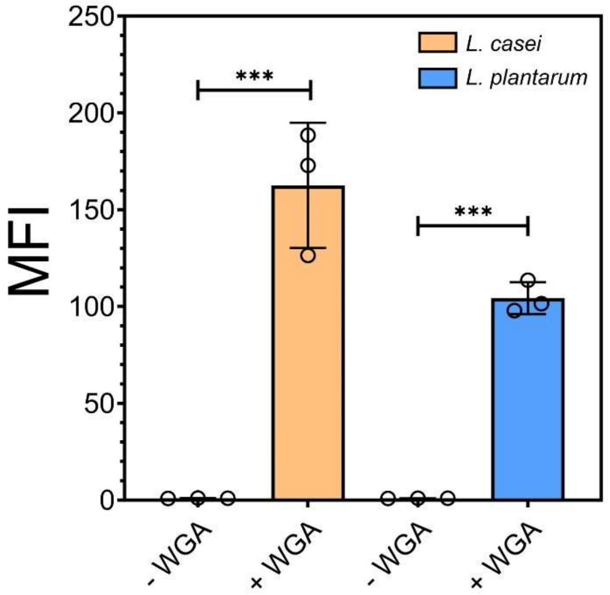


**Figure S3.** Flow cytometry analysis of *L. casei* and *L. plantarum* sacculi in the presence of DMSO or WGA-fluorescein (1 μM) for 60 min then washed with 1X PBS. Mean fluorescence intensity (MFI) is the ratio of fluorescence levels above the control (DMSO) treatment from 10000 events. P-values were determined by a two-tailed t-test (* denotes a p-value < 0.05, ** < 0.01, ***<0.001, ns = not significant).

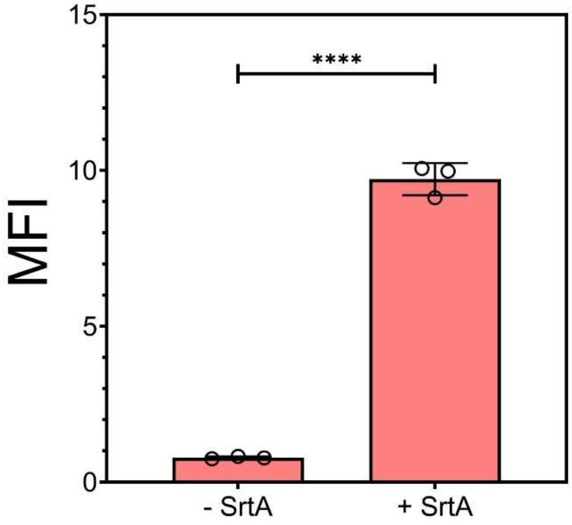


**Figure S4.** Flow cytometry analysis of sacculi isolated from *L. casei* incubated with 100 μM Fl-LMPTG (substrate) in the presence or absence of SrtA (20 μM) for 5 h then washed with 100 mM Tris, 5 mM EDTA, pH 7 and freshly added 8 M urea. Mean fluorescence intensity (MFI) is the ratio of fluorescence levels above the control (-SrtA) treatment from 10000 events. P-values were determined by a two-tailed t-test (* denotes a p-value < 0.05, ** < 0.01, ***<0.001, ns = not significant).

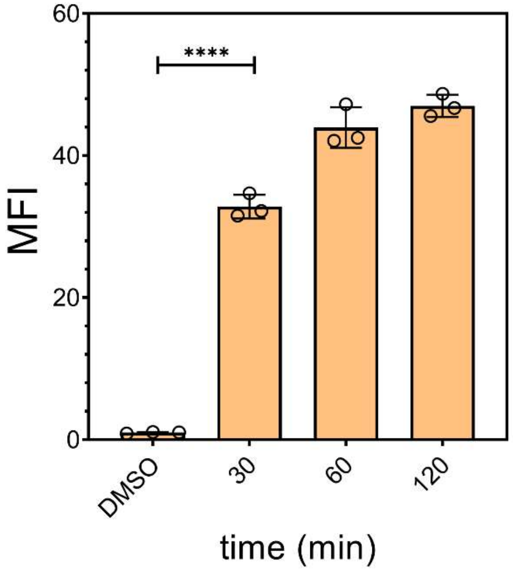


**Figure S5**. Flow cytometry analysis of sacculi of *L. casei* incubated with **DAzDA** overtime. First, *L. casei* cells were incubated with 1 mM of DAzDA or DMSO (control). *L. casei* cells samples were collected 30, 60, and 120 min after supplementation of the probe. Next, samples from the different time point where subjected to sacculi isolation then treated with 30 μM of alkyne-fluorescein. Mean fluorescence intensity (MFI) is the ratio of fluorescence levels above the control (DMSO) treatment from 10000 events. P-values were determined by a two-tailed t-test (* denotes a p-value < 0.05, ** < 0.01, ***<0.001, ns = not significant).

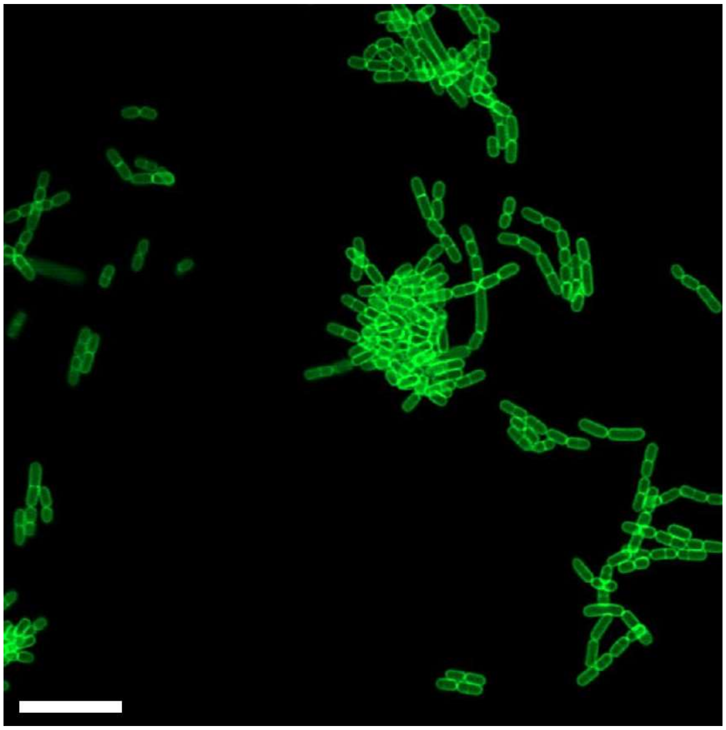

**Figure S6.** Confocal microscopy analysis of *L. casei* sacculi labeled with 1 mM **DAzDA** followed by subsequent click chemistry (CuAAC) of 30 μM Fluorescein-alkyne for 1 h at 37°C; scale bar = 5 μm.

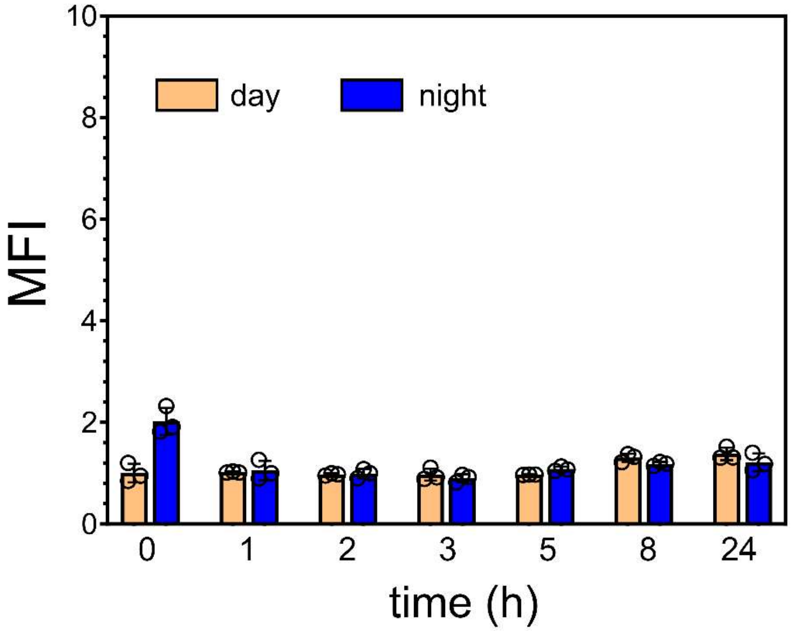


**Figure S7.** SPF mice were orally dosed with 5 mM of **DAzDA** 2X one hour apart. Following the second dosing, the animals were moved to a new cage. Fecal samples were collected at the designated times and subjected to sacculi isolation and a click reaction to install the fluorophore (in the absence of the copper catalyst). The process was performed during daylight and during the night cycles. Mean fluorescence intensity (MFI) is the ratio of fluorescence levels above the control (DMSO) treatment from 10000 events.

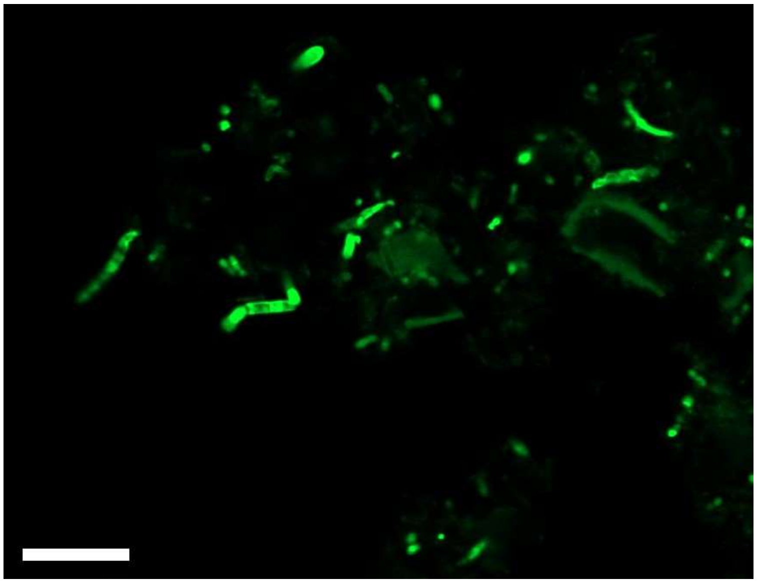

**Figure S8.** Confocal microscopy of gut bacterial sacculi labeled with 5 mM **DAzDA** followed by subsequent click chemistry (CuAAC) of 30 μM Fluorescein-alkyne for 1 h at 37 °C; scale bar = 5 μm.

**Materials.** Starting materials including protected amino acids and reagents needed for peptide synthesis were purchased from Chem-Impex. Vancomycin-BODIPY (Catalog # V34850) was purchased from ThermoFisher. Wheat-germ albumin-Fluorescein (WGA-FI) (catalog # FL-1021) was purchased from vector laboratories. All other organic chemical reagents were purchased from Fisher Scientific or Sigma Aldrich and used without further purification.

**Mice**. Wild type mice were obtained from The Jackson Laboratory. All experiments were conducted utilizing adult, female mice bred and maintained in pathogen-free barrier facilities at the University of Virginia. All animal experiments in this study were approved by the University of Virginia Institutional Animal Care and Use Committee.

**Flow cytometry analysis of whole bacterial cells and isolated peptidoglycan/ sacculi.** Bacteria or isolated peptidoglycan from experiments below were analyzed using an Attune NxT flow cytometer equipped with a 488 nm laser and 525/40 nm bandpass filter. The data were analyzed using the Attune NxT Software, where populations were gated and no less than 10,000 events per sample were recorded. Wherever presented the mean fluorescence intensity (MFI) is the ratio of fluorescence levels above the control treatment.

**Isolation of *L.casei* and *L.plantarum* peptidoglycan/ sacculi.** Methodology for isolation of sacculi is an adaptation from published protocols.[1] *L. casei*/ *L plantarum* bacteria were added to MRS broth (1:100) and allowed to grow overnight at 37°C with shaking at 250 rpm. The cells were harvested at 10,000g for 10 min at 4°C. The resulting pellet was resuspended in 3 mL of 1X PBS containing 4% SDS and boiled at 100°C for 1 h. The samples were washed 6 times with Milli-Q water to remove any residual SDS from the previous treatment. Then the samples were resuspended in 0.1M Tris pH 8 and subjected to stepwise digestion using 15 μg/mL DNAse / 60 μg/mL RNAse and 50 μg/mL trypsin at 37°C for 1 h, respectively. The samples were boiled for 5 min at 100°C to inactivate the enzymes, followed by washing twice with Milli-Q water. The isolated peptidoglycan was then resuspended in 1M HCL and incubated for 4 h at 37°C to remove the wall teichoic acids. The sacculi were recovered by centrifugation at 21,000*g* for 20 min, washed 6X with 0.1 M tris pH 8, resuspended in Milli-Q water, and stored at –20°C for subsequent analysis.

**Isolation of gut commensal peptidoglycan/ sacculi from mice and human fecal samples.** Fecal sample from a single healthy human donor was purchased from lee Biosolutions (catalog # 991-18). Fecal pellets (mice or human) were resuspended in sterile 1X PBS, homogenized by sonification and vortexing for 30 min. The fecal pellets suspensions were strained using a 40 μm cells strainer (pluriselect catalog # 43-10040-50) to get rid of debris. The bacterial cells were obtained by centrifugation 21,000g for 20 min. The bacterial cells were then boiled in 4% SDS at 100°C for 1 h. The samples were washed 6X with Milli-Q water to remove any residual SDS from the previous treatment. Then, the samples were resuspended in 0.1M Tris pH 8 and subjected to stepwise digestion using 15 μg/mL DNAse / 60 μg/mL RNAse and 50 μg/mL trypsin at 37°C for 1 h, respectively. The solutions were boiled for 5 min at 100°C to inactivate the enzymes, followed by washing twice with Milli-Q water. The isolated sacculi was then resuspended in 1M HCL and incubated for 4 h to remove the wall teichoic acids. The sacculi were recovered by centrifugation at 21,000g for 20 min, washed 6X with 0.1 M tris pH 8, resuspended in Milli-Q water and stored at –20°C for subsequent analysis.

**Vancomycin-BODIPY (VBD) binding and competition Assays.** *L. casei* sacculi or gut bacterial/ fecal sacculi samples were pelleted in a 96-well plate at 3300g for 4 min and then resuspended in a 1X PBS solution containing 2 μg/mL VBD. The plates were incubated for 1 h at 37°C, washed 3 times with 1X PBS, and analyzed by flow cytometry as described above. For competition of VBD, the fecal sacculi were resuspended in 1X PBS solution containing 2 μg/mL VBD or 2 μg/mL VBD in conjunction with either 50, 100, 500, 1000, or 10000 μg/mLof the L-Lys-D-Ala-D-Ala peptide. The plates were incubated for 1 h at 37°C, then washed 3X with 1X PBS and immediately subjected to analysis by flow cytometry as described above**.**

**LysM-GFP expression and purification.** LysMD3-GFP cDNA was synthesized by BioMatik on a pET-21a(+) vector with ampicillin resistance gene. Competent BL21 cells (DE3, gold, Agilent, cat # 230130) were transformed with the plasmid. A single colony of cells grown on ampicillin agar plate was picked up and resuspended in LB media (5 mL) and incubated at 37°C for 1 h. The media was then added to 2 L fresh LB media supplemented with ampicillin for 6 h at 37°C. Upon addition of 1M Isopropyl-β-D-thiogalactopyranoside (IPTG), the temperature of incubator was lowered to 18°C and incubated for 18 h. The cells were collected by centrifugation at 4000g for 30 min and resuspended in buffer (50 mM Tris, 300 mM NaCl, pH 8.0) in an ice bath.[2] To this suspension of cells, 2.5 mg of lysozyme was added (MP Biomedicals LLP, Cat# 100834, Lot #MR28337) and 2 tablets of complete Mini protease inhibitor (Roche Diagnostics, Lot#53945000). Cells were lysed using an ultrasonic cell sonicator and the lysate was separated from cellular debris by centrifugation at 20,000g for 30 min at 4°C. The supernatant was loaded onto a Ni-NTA agarose resin packed column and eluted with 250 mM imidazole in buffer (50 mM Tris, 300 mM NaCl, pH 8.0). The eluted protein was dialyzed against buffer without imidazole, aliquoted (protein concentration determined by Bradford assay), and analyzed by gel electrophoresis and fluorescence spectroscopy and stored at –20°C until use. Protein sequence:

MVSKGEELFTGVVPILVELDGDVNGHKFSVSGEGEGDATYGKLTLKFICTTGKLPVPW
PTLVTTLTYGVQCFSRYPDHMKQHDFFKSAMPEGYVQERTIFFKDDGNYKTRAEVKF
EGDTLVNRIELKGIDFKEDGNILGHKLEYNYNSHNVYIMADKQKNGIKVNFKIRHNIEDG
SVQLADHYQQNTPIGDGPVLLPDNHYLSTQSALSKDPNEKRDHMVLLEFVTAAGITLG
MDELYKGGGGSGGGGSIVLTKDIQEGDTLNAIALQYCCTVADIKRVNNLISDQDFFALR
SIKIVLEHHHHHH (305 amino acids, MW: 34 kD)

**LysM-GFP and WGA-Fl sacculi binding assay.** Isolated Fecal sacculi samples, resuspended in PBS, were incubated with either 1 μM LysM-GFP,1 μM WGA-Fl or no protein. The plates were incubated for 1 h at 37°C, then washed 3X with 1X PBS, and analyzed by flow cytometry as described above.

**Sortase A Expression and Purification.** The plasmid for sortase A from *S. aureus* was obtained from Addgene: pET28a-SrtAdelta59. Competent BL21 (DE3) *E. coli* cells were transformed with the plasmid. *E. coli* cells containing the plasmid from an overnight culture were diluted 1:100 and grown at 37°C until the $OD_{600}$ was 0.4-0.6. At this time, Isopropyl-β-D-thiogalactopyranoside (IPTG) was added to a concentration of 0.5 mM and the cultures were shaken at 25°C for 18 h. Cells were collected at 3,000g for 30 min and resuspended in 50 mM Tris-HCl, pH 7.5, and 150 mM NaCl. Pellets were then recollected and lysed in cold lysis buffer (50 mM Tris-HCl, pH 7.5, 150 mM NaCl, 5 mM $MgCl_2$, 10 mM imidazole, 10% vol/vol glycerol, 1 mg $ml^{-1}$ DNase and 1 mg $ml^{-1}$ lysozyme). Cells were lysed using an ultrasonic cell sonicator and the lysate was separated from cellular debris by centrifugation at 20,000g for 30 min at 4°C. The supernatant was loaded onto a Ni-NTA agarose resin and eluted with 500 mM imidazole in buffer (50 mM Tris-HCl pH 7.5, 150 mM NaCl, and 10% vol/vol glycerol). The eluted protein was dialyzed against buffer without imidazole, aliquoted (protein concentration determined by Bradford assay), and stored at –20°C until use.
**Sortase A Enzymatic Assays.** Isolated fecal sacculi samples were incubated with 20 μM sortase A or no enzyme, 100 μM sorting signal substrate (Fl-LPMTG), and 1X sortase buffer (10X contains 500 mM TrisHCl, pH 7.5, 1.5 M NaCl, 100 mM $CaCl_2$). The samples were shaken at room temperature for 5 h. The enzymatic activity was quenched with 0.1% TFA, and washed 3X with 100 mM Tris, 5 mM EDTA, pH 7 and freshly added 8 M urea. Samples were resuspended in a final volume of 200 μL 1X PBS and analyzed by flow cytometry.

**Enzymatic degradation of fecal sacculi. Fl-LPMTG** (100 μM) labeled fecal sacculi samples were pelleted in a 96-well plate at 3300g for 4 min and then resuspended in a 1X PBS solution containing 25 μg/mL mutanolysin and allowed to incubate at 37°C. A zero time point was taken before any samples were subjected to enzyme treatment. A portion of the sacculi were then taken at 1, 5, 15, 30, and 60 min. At each time point, the collected sacculi were resuspended in a solution of 1X PBS containing 4% formaldehyde to quench the enzymatic activity. Samples were analyzed by flow cytometry.

**Peptidoglycan processing.** Intact sacculi were resuspended in 400μL of final concentration 5μM NaHPO4/NaH2PO4 buffer. Samples were digested with 3000U of mutanolysin shaking overnight at 37°C. After initial digestion, an additional 3000U of mutanolysin was added and digested for 4 hours before enzyme inactivation (100 °C for 15 miin). Undigested material was pelleted by centrifugation (20,000g for 45 min), the muropeptide-containing supernatant was frozen, and dried using a high vacuum line coupled with a liquid nitrogen solvent trap. The dried muropeptides were resuspended in 300 μl of saturated sodium borate buffer, pH 9.25, which was aliquoted into two equal volumes. 50 mg of sodium borohydride or deuteride was dissolved in 500 μl of LCMS grade water and 50 μl of either was slowly added to separate muropeptide aliquots in a dropwise manner. Approximately 10 μl of formic acid was added after 1 h to quench the reaction and adjust the pH to 3. The samples were snap frozen and dried once more with a high vacuum line and stored in a desiccator until analysis, at which time the samples were resuspended in H2O: MeCN (200 μl, 9:1, v:v) containing 0.1% formic acid. The samples were sonicated in a water bath for 10 min and centrifuged at 13,000 x g for 10 min at 4 °C. 180 µl of the supernatant was transferred to an LCMS vial for analysis.

**LCMS analysis of muramidase digested peptidoglycan.** Analysis was performed on a Shimadzu LCMS9030 QToF system coupled to a LC-40B X3 UPLC, a SIL-40C X3 autosampler (10 °C), and a CTO-40C column oven (40 °C). Reduced mutanolysin-digested muropeptides were separated on a Waters BEH C18 column (2.1x100 mm, 130 Å pore size, 1.7 µm particle size), using a binary gradient separation at a flow rate of 300 uL/min. Solvent A was water and Solvent B was MeOH, with both solvents containing 0.1% formic acid (v/v). The initial solvent condition was 99:1 (A: B, 0-3 min). Stepwise linear gradients first to 8% Solvent B (3-12 min), then to 20% Solvent B (12-24 min), and finally 5% Solvent B (24-25 min), was followed by a hold at 5% B until 30 min. The column was then re-equilibrated to 1% B (1 min to 1%B followed by a 5 min re-equilibration time). The initial 1.25 min of each separation was diverted to waste to prevent contamination of the mass spectrometer interface with salts from the reduction step. The samples were subjected to electrospray ionization in positive ion mode using data dependent acquisition. The interface voltage was 4.0 kV at 300 °C; desolvation temperature of 526 °C; DL temperature of 250 °C. Nebulizing, heating, and drying gas flows were at 2, 10, and 10 L/min, respectively. The m/z range for the MS scan was 300 to 2000 with fragmentation data collected from 50-2000 m/z with a collision energy of 30 (+/- 5). Muropeptides were identified based upon ions that provided known muropeptide masses and were fragmented to release both the GlcNAc moiety (204.085 m/z) and a GlcNAc ring cleavage product (138.053 m/z).

**Metabolic labeling of *L. casei* and sacculi isolation.** MRS (3 mL) containing 1 mM **DKAz,** 1mM **DAzDA** or no probe was prepared. *L. casei* cells were added to the MRS broth (1:100) and allowed to grow overnight at 37°C with shaking at 250 rpm. Cells were then harvested at 10,000g for 10 min at 4°C and washed 3X in 1X PBS and resuspended in 1X PBS containing 4% SDS and boiled for 1hr. The samples were washed 6X with Milli-Q water to remove any residual SDS residue from the previous treatment. Then the samples were resuspended in 0.1M Tris pH 8 and subjected to stepwise digestion using 15 µg/mL DNAse /60 µg/mL RNAse and 50 µg/mL trypsin at 37°C for 1 h, respectively. The solutions were boiled for 5 min at 100°C to inactivate the enzymes, followed by washing twice with Milli-Q water. The isolated sacculi was then resuspended in 1M HCL and incubated for 4 h to remove the wall teichoic acids. The sacculi were recovered by centrifugation at 21,000g for 20 min, washed 5 X with 0.1 M tris pH 8, resuspended in Milli-Q water and stored at -20°C for subsequent analysis.

**Time dependent Labeling of *L. casei* sacculi with DAzDA.** *L. casei* bacteria were added to the MRS broth (1:100) and allowed to grow overnight at 37°C with shaking at 250 rpm to an Optical density (OD) of 1.0. Then, the cells were incubated with 1mM of DAzDA for 30, 60, or 120 min. Then, *L. casei* sacculi was isolated as previously described. Then, the incorporation of the probe into the peptidoglycan was monitored by CuAAC "click chemistry" of Fl-alkyne (30 µM) as described below. The fluorescent sacculi was analyzed by flow cytometry as described above.

***In-vivo* gut labeling with DKAz, DAzDA or DAzLA.** Female Mice (16 -18 weeks old) were orally gavaged twice with 200 µL of 5 mM of **DKAz**, **DAzDA** or **DAzLA**, one hour apart during their day metabolism (light) cycle. Then, fecal pellets were collected 1, 2, 3, 5, 8, and 24 h post-gavage. Collected fecal pellets were immediately frozen at -80°C for later subsequent sacculi isolation and analysis. The same experiment was repeated 3 days later during the night metabolism (dark) cycle of the mice to evaluate changes in labeling trends. Isolation of sacculi was undergone as described previously.

**CuAAC/ click reaction on bacterial peptidoglycan/ sacculi.** Sacculi samples (10 µL) were pelleted in a 96-well plate at 3300g for 4 min and then resuspended in a 1X PBS solution containing 1 mM CuSO4, 128 µM THPTA, 1.2 mM freshly prepared sodium ascorbate, and 30 µM of Fluorescein-alkyne (Lumiprobe catalog # C41B0) in PBS. The reaction components were added in the exact order listed, with sodium ascorbate and alkyne-fluorescein added immediately before incubation. CuAAC reactions were performed at 37°C for 60 min. The sacculi were then washed 3X with 1X PBS and analyzed by Flow cytometry as described above.

**Confocal imaging of labelled sacculi.** Glass microscope slides were spotted with 1% agarose pads and 2 µL of labeled bacterial/ sacculi samples were deposited onto each agarose pad. Samples were covered with cover slips and imaged using a Zeiss 880/990 multiphoton Airyscan microscopy system (zoomed with a 63x oil-immersion lens) equipped with a 488 nm laser. Images were obtained and analyzed via Zeiss Zen software. We acknowledge the Keck Center for Cellular Imaging and for the usage of the Zeiss 880/980 multiphoton Airyscan microscopy system (PI- AP: NIH-OD025156).

#### Synthesis

#### Scheme S1. synthesis of DAzDA and DAzLA

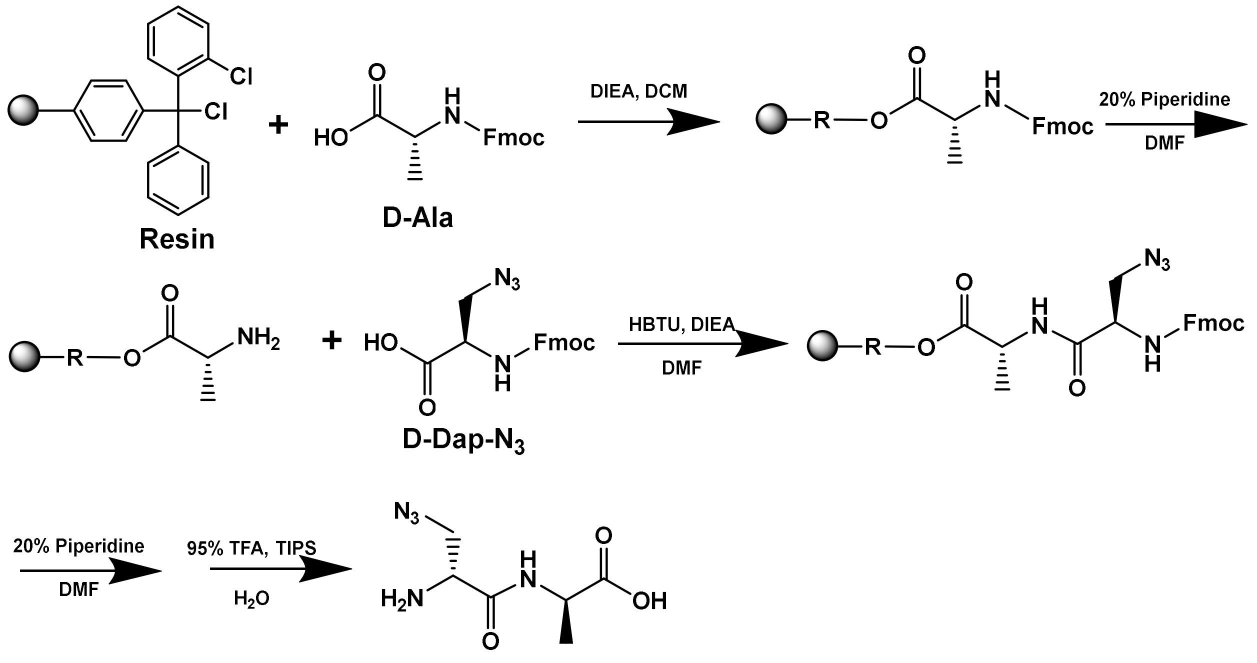


**D-Dap-$N_3$-D-Ala**

To a 25 mL peptide synthesis vessel with 200 mg 2-Chlorotrityl chloride resin (0.142 mmol) resuspended in 15 mL dry dichloromethane, was added Fmoc-D-alanine or Fmoc-L-alanine (97 mg,1.1 eq, 0.3124 mmol), and diisopropylethylamine (DIEA, 4.4 eq, 0.11 mL, 1.2496 mmol). The resin was shaken for 1 h at room temperature and washed with methanol and dichloromethane (3 x 15 mL each alternatively). The Fmoc protecting group was removed with a 20% piperidine in DMF solution (15 mL) for 30 min at ambient temperature, then washed with MeOH and DCM (3 x 15 mL each). Boc-D-DapN3.CHA was used for the next coupling. To remove the cyclohexylammonium salt, 3 equivalent of the amino acid was dissolved in 20 mL of ethyl acetate in a separatory funnel. The compound was washed twice with 1M HCL ( total 20 mL), then with a saturated NaCl solution (total 20 mL), and finally rotary evaporated to dryness. For coupling, the amino acid was dissolved in DMF (15 mL) containing HBTU (3.0 eq, 0.323 g, 0.852 mmol), and DIEA (6.0 eq, 0.297 mL, 1.704 mmol) in DMF (15 mL) was added to the vessel and shaken for 2 h at room temperature. The resin was washed as previously described. To remove the peptide from resin, a TFA cocktail solution (95 % TFA, 2.5% TIPS, and 2.5 % DCM) was added to the resin with agitation for 2 h protected from light. The resin was filtered, and the resulting solution was concentrated *in vacuo*. The peptide was triturated with cold diethyl ether.

**Scheme S2. Synthesis of Fl-LPMTG**

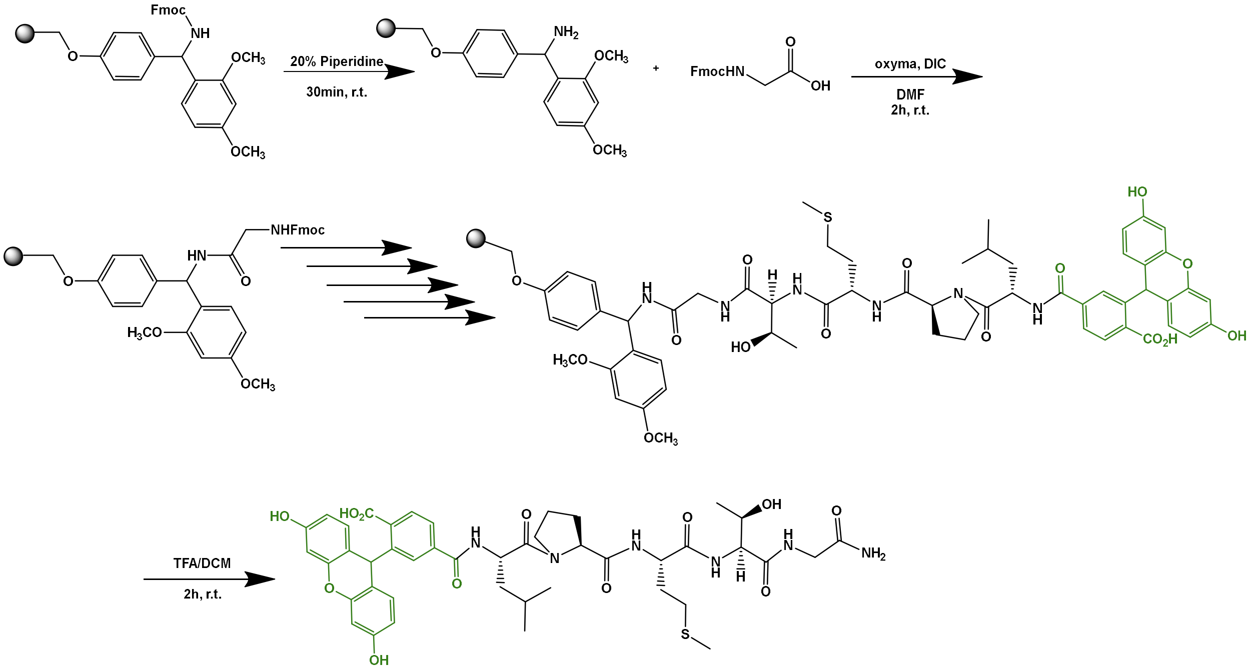


A 25 mL synthetic vessel was charged with 500 mg (0.27 mmol) of Fmoc-Rink amide resin. The Fmoc protecting group was removed with a 20% piperidine in DMF solution (15 mL) for 30 min at ambient temperature, then washed with MeOH and DCM (3 x 15 mL each). Fmoc-glycine-OH (3 eq, 240 mg, 0.810 mmol), Oxyma (3 eq, 115 mg, 0.810 mmol), and DIC (3 eq, 0.126 mL, 0.810 mmol) in DMF (15 mL) was added to the reaction vessel and agitated for 2 h at ambient temperature and washed as previously stated. The Fmoc removal and coupling procedure was repeated as before using the same equivalencies with Fmoc-L-threonine(tBu)-OH, Fmoc-Lmethionine-OH, Fmoc-L-proline-OH, and Fmoc-L-leucine-OH. The Fmoc group of L-leucine was deprotected and resin coupled with 5,6-carboxyfluorescein (2 eq, 203 mg, 0.810 mmol), HBTU (2 eq, 201 mg, 0.810 mmol), and DIEA (4 eq, 0.187 mL, 1.08 mmol) in DMF (15 mL) shaking overnight. The resin was washed as previously described. To remove the peptide from resin, a TFA cocktail solution (95 % TFA, 2.5% TIPS, and 2.5 % DCM) was added to the resin with agitation for 2 h protected from light. The resin was filtered, and the resulting solution was concentrated *in vacuo*. The peptide was triturated with cold diethyl ether yielding the crude peptide. The peptide was purified using reverse phase HPLC (RP-HPLC) using a Waters 1525 Binary HPLC Pump using a Phenomenex Luna 5u C8(2) 100A (250 x 4.60 mm) prep column with an eluent consisting of solvent A ($H_2O$ / 0.1% TFA) and solvent B (MeOH / 0.1% TFA) using a 60 min gradient transitioning from 5% B to 100% B at a flow rate of 1.5ml/min to yield **Fl-LPMTG**. The purity of the peptides was verified by analytical RPHPLC using a Phenomenex C8 column with an eluent consisting of solvent A ($H_2O$ /0.1% TFA) and solvent B (MeOH / 0.1% TFA) with a 30 min gradient transitioning from 5% B to 100% B at a flow rate of 1.9 ml/min and monitored at 220 nm and molecular weight was confirmed using MALDI-TOF [M + $H^+$]: 875.3280.

**Compound characterization.**

**High Resolution Mass Spectrometry(HRMS) and NMR analysis of DAzDA and DAzLA dipeptides.** High resolution mass spectrometry (HRMS) analysis of the compounds was done using a C18(2) column (Luna 5 µm 100Å 250 x 4.6 mm) connected to an Agilent LC-QTOF (Agilent 1260 Infinity II Prime LC with Agilent 6545B QTOF). The mass of the peptides was analyzed by MS using MassHunter software. Further analysis of purity and characterization of dipeptides was done through $^{1}$H NMR.

**ESI-MS calculated [M + H+]: 202.0935, found: 202.0968**

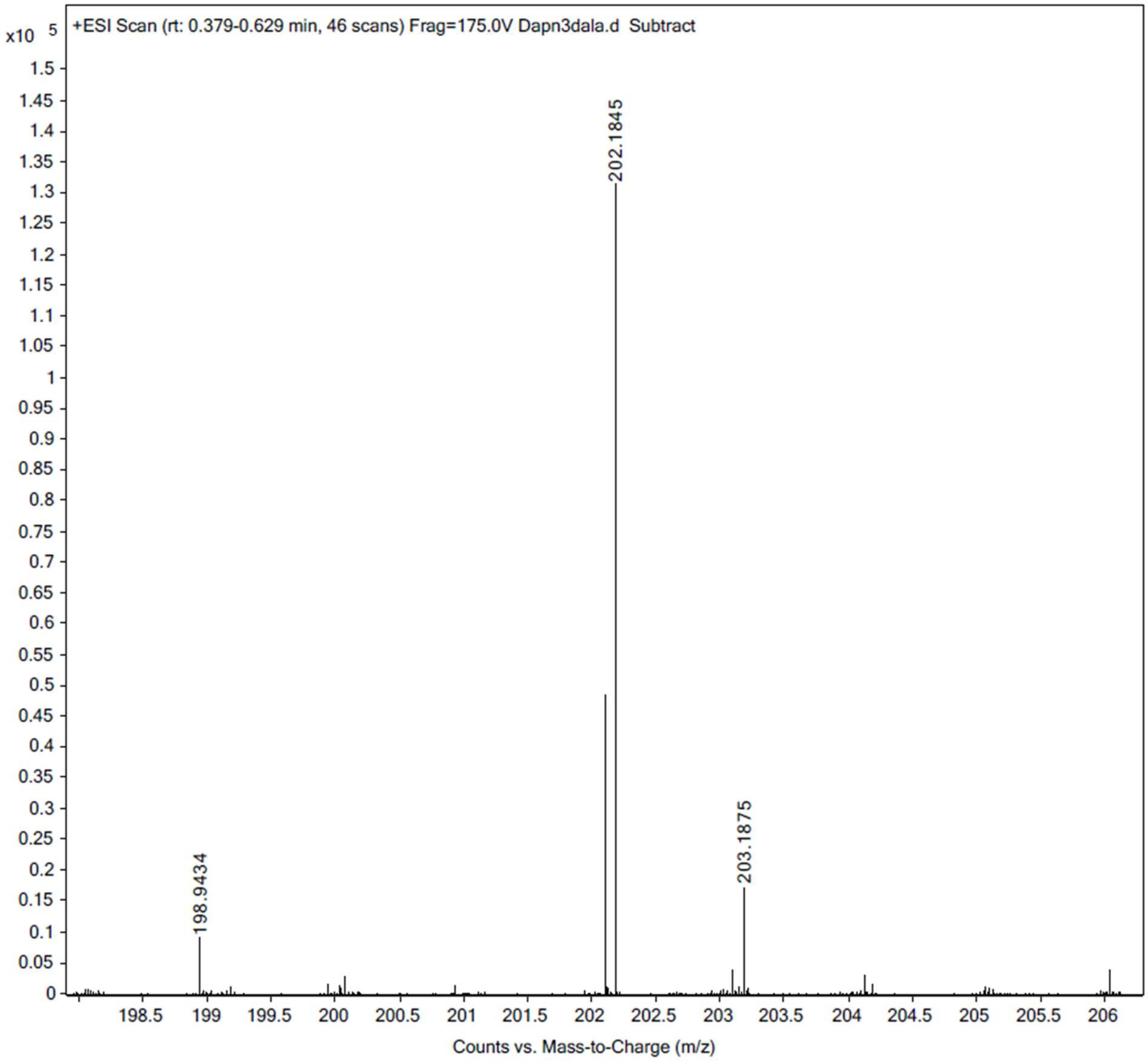

**D-dipeptide.** $^{1}H$ NMR (dmso-d6) d 1.31 (d, *J* = 6.0 Hz, 3H, $CH_3$), 3.66 (dd, *J* = 6.0 and 12.0 Hz, 1H, $CH_2$), 3.82 (dd, *J* = 6.0 and 12.0 Hz, 1H, $CH_2$), 4.02 (dd, *J* = 6.0 and 12.0 Hz, 1H, $CH_2$), 4.27 (m, *J* = 6.0 and 12.0 Hz, 1H, CH), 8.87 (brd, *J* = 6.0 Hz, 2H, $NH_2$).

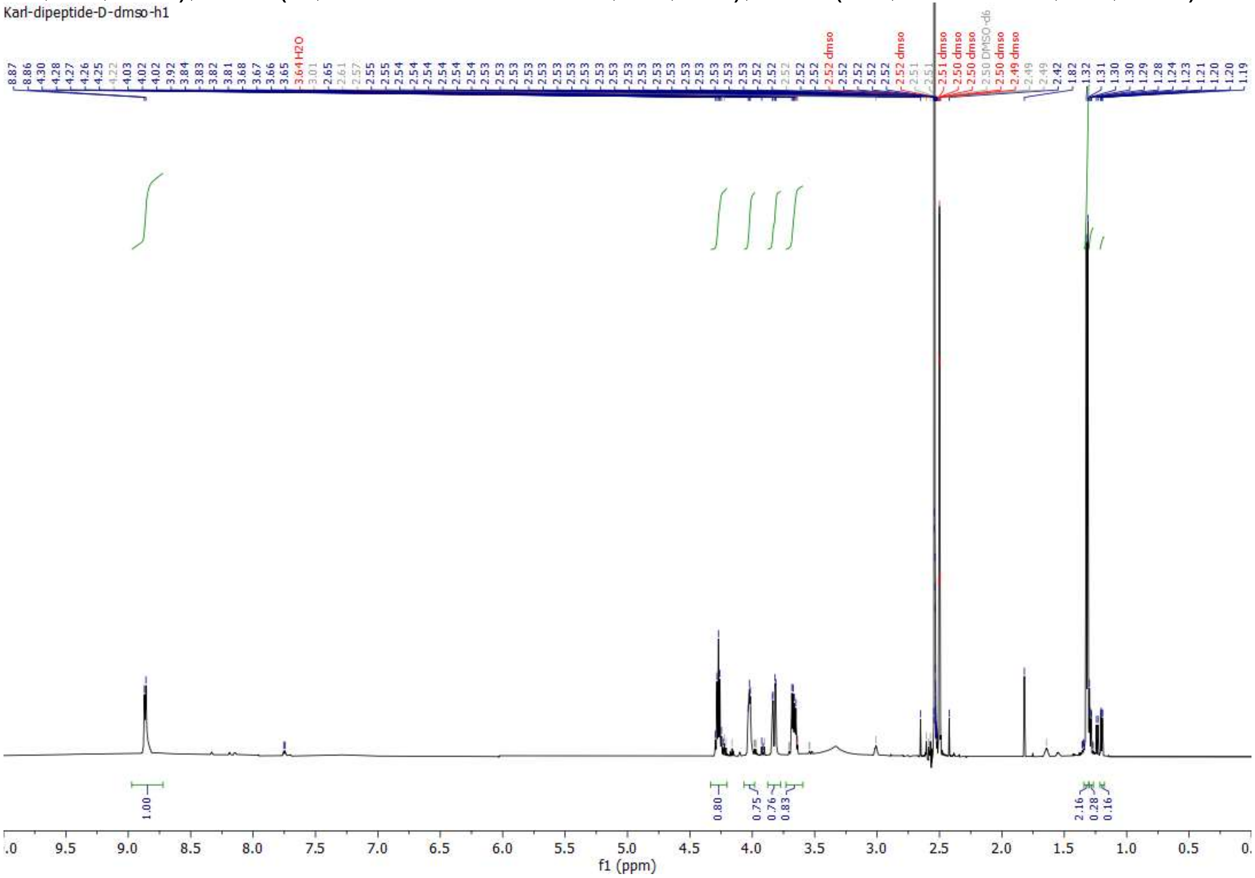

**ESI-MS calculated [M + H+]: 202.0935 , found: 202.0968**

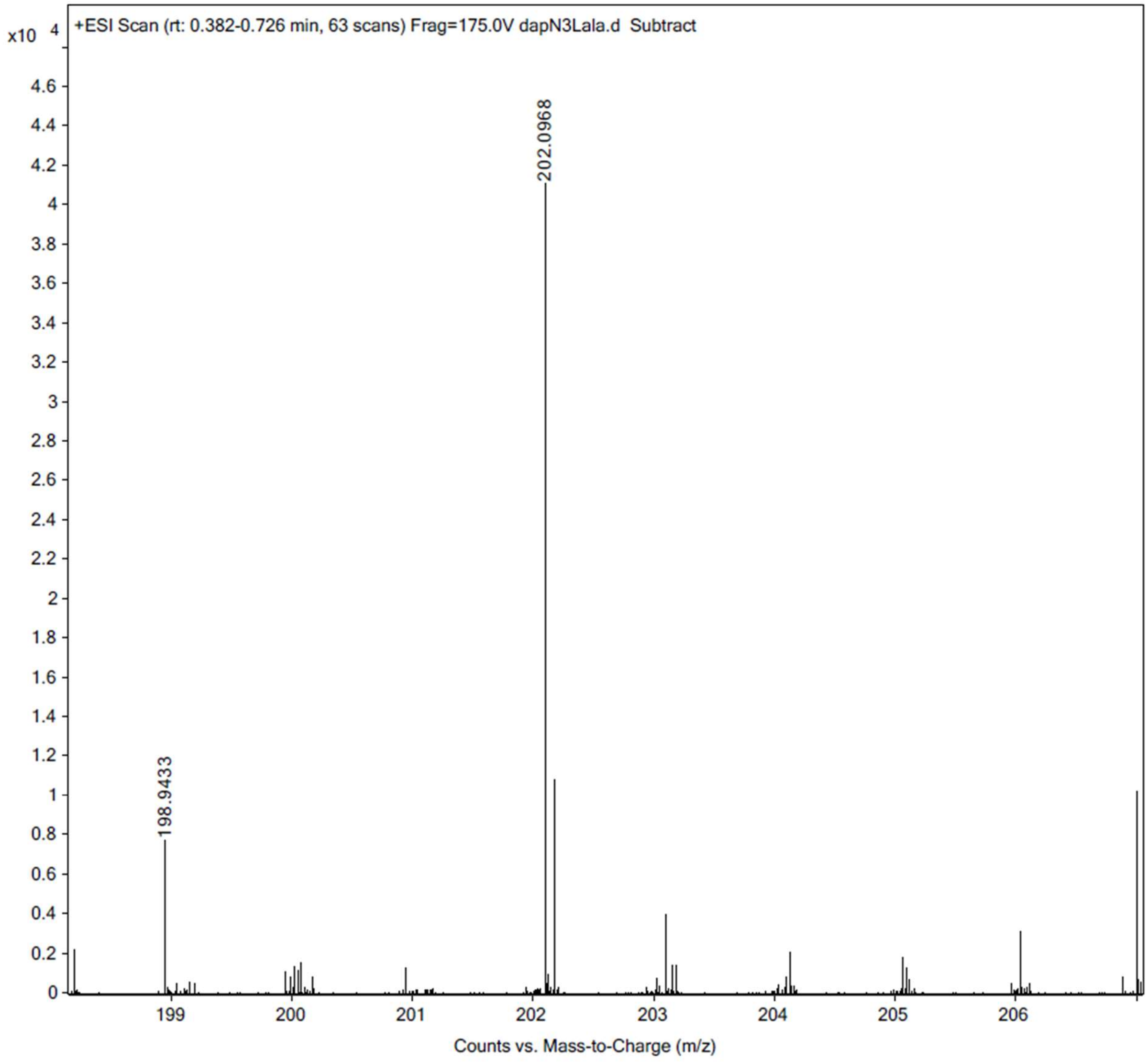

**L-Dipeptide.** $^1$H NMR (dmso-d6) d 1.30 (d, *J* = 6.0 Hz, 3H, $CH_3$), 3.66 (dd, *J* = 6.0 and 12.0 Hz, 1H, $CH_2$), 3.76 (brd, *J* = 6.0 and 12.0 Hz, 2H, $CH_2$), 4.04 (dd, *J* = 6.0 and 12.0 Hz, 1H, $CH_2$), 4.28 (m, *J* = 6.0 and 12.0 Hz, 1H, CH), 8.88 (brd, *J* = 6.0 Hz, 2H, $NH_2$).

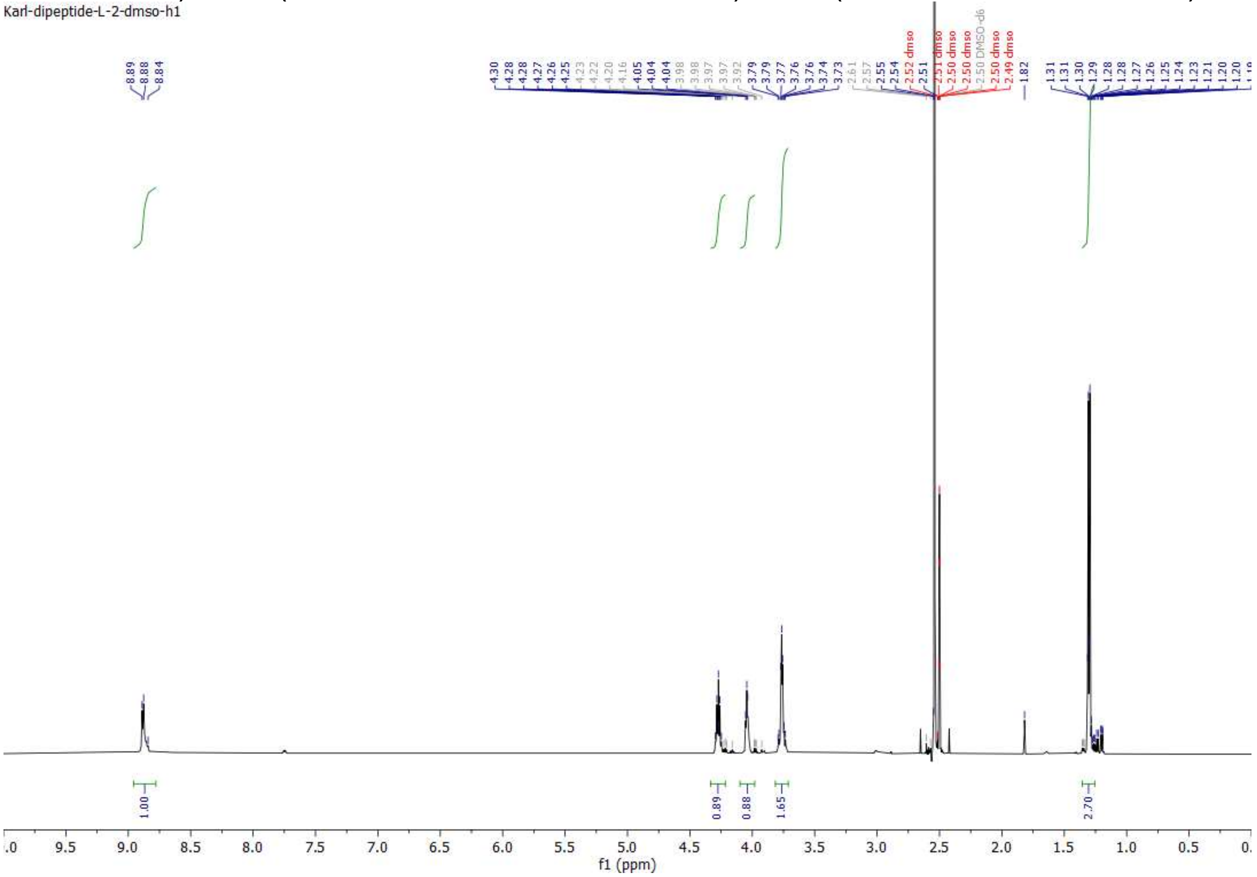

**Fl-LPMTG analytical HPLC.**

**Fl-LPMTG** was analyzed for purity using a Waters 1525 Binary HPLC Pump using a Phenomenex Luna 5u C8(2) 100A (250 x 4.60 mm) column; gradient elution with H2O/CH3OH + 0.1% TFA.The large peak indicate the DMSO frontline.

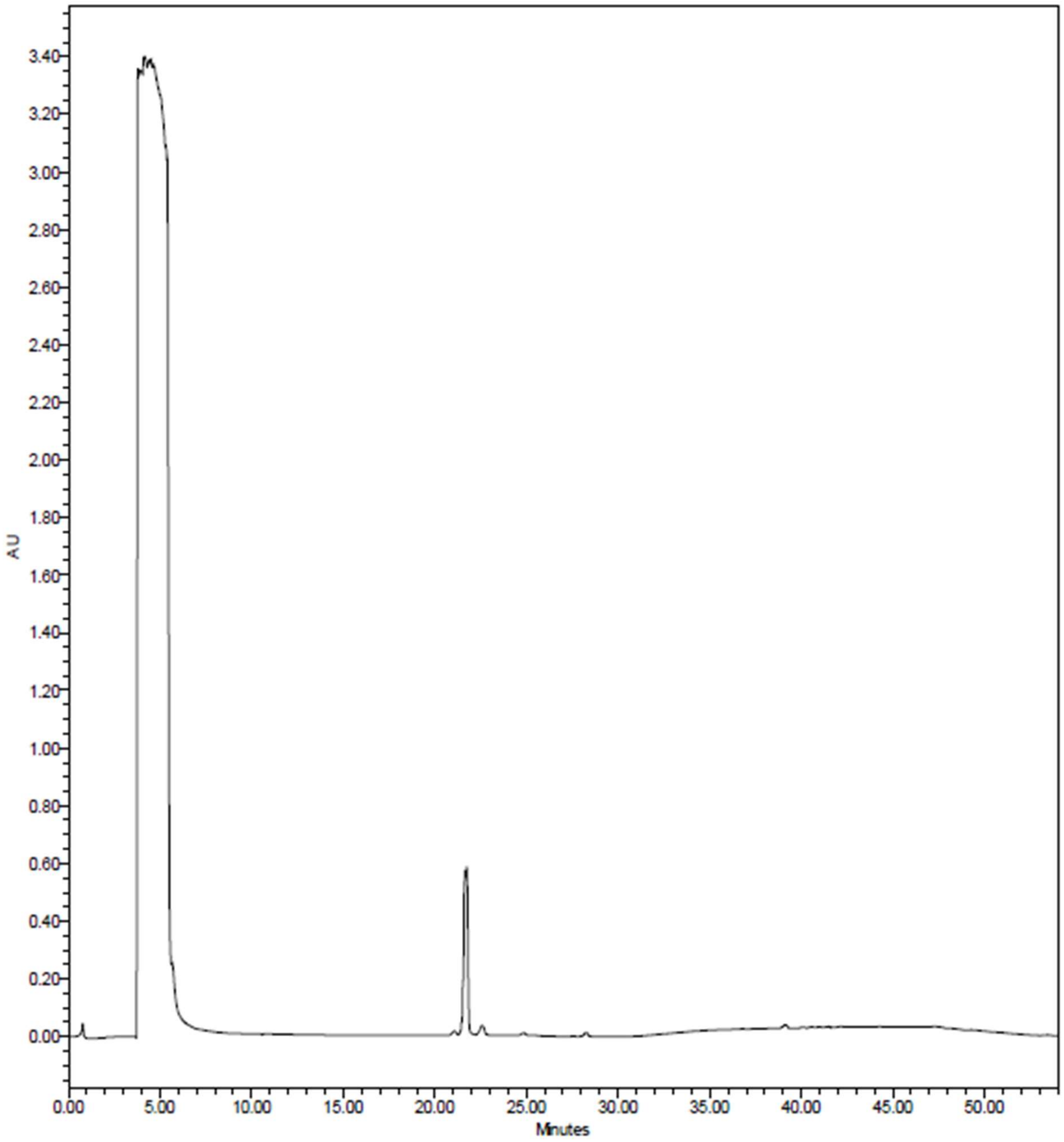

**Fl-LPMTG High Resolution Mass spectrometry (HRMS) analysis.**
**ESI-MS calculated [M + H+]: 875.3280, found: 875.3242**

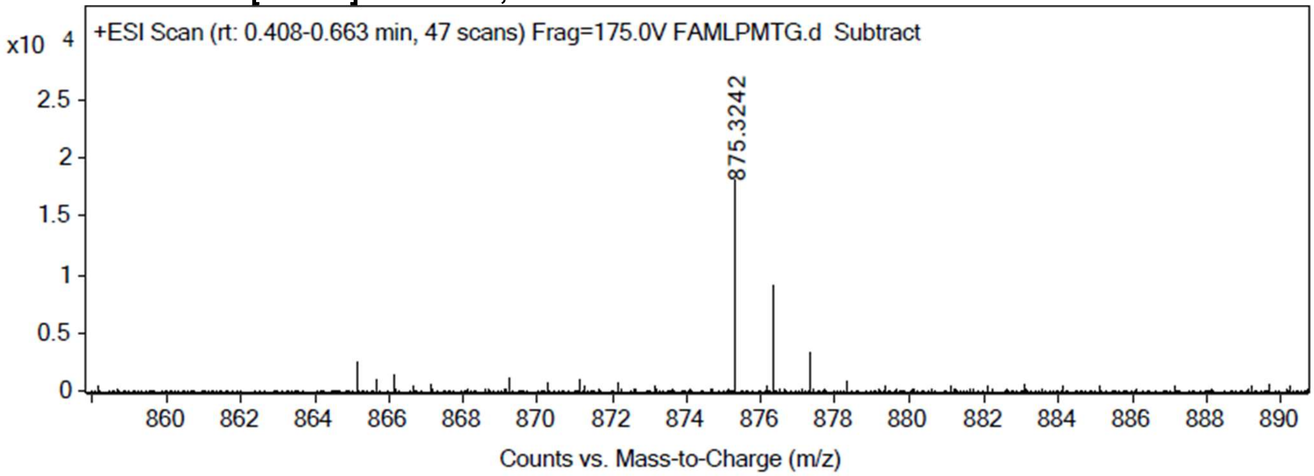

#### References

1. Kuhner, D.; Stahl, M.; Demircioglu, D. D.; Bertsche, U., From cells to muropeptide structures in 24 h: peptidoglycan mapping by UPLC-MS. *Sci Rep* **2014,** *4*, 7494.
2. Mesnage, S.; Dellarole, M.; Baxter, N. J.; Rouget, J. B.; Dimitrov, J. D.; Wang, N.; Fujimoto, Y.; Hounslow, A. M.; Lacroix-Desmazes, S.; Fukase, K.; Foster, S. J.; Williamson, M. P., Molecular basis for bacterial peptidoglycan recognition by LysM domains. *Nat Commun* **2014,** *5*, 4269.
